# Laminin-511 protects pancreatic β-cells from cytokine-induced death through integrin-mediated pro-survival signaling and modulation of protein kinase C δ

**DOI:** 10.64898/2026.08.04.742875

**Authors:** Christine El-Dirani, Meghana Hosahalli Shivananda Murthy, Keifer Holcomb, Gwendolyn Gutierrez, Riley Starzel, Brisa Peña, Daewon Park, Richard K.P. Benninger, Nikki L. Farnsworth

## Abstract

During the progression of type 1 diabetes (T1D), the extracellular matrix (ECM) surrounding pancreatic islets is degraded concurrent with infiltration of autoreactive immune cells and β-cell death. Among the lost ECM proteins, laminin-511 is known to be essential for islet survival under healthy and T1D associated conditions, including high levels of pro-inflammatory cytokines. However, the key β-cell signaling pathways regulated by laminin and the contributions to T1D pathogenesis when these cues are lost are poorly understood. This study utilizes a biomimetic reverse thermal gel (RTG) with laminin-511 to determine if laminin protects β-cells against cytokine-induced death and elucidate the signaling pathways involved. MIN6 cells, C57Bl/6 mouse islets and human islets were encapsulated in RTG scaffolds with laminin-511 and treated with a cytokine cocktail for 24 hours. Islet viability and the activities of several pro- and anti-apoptotic proteins were studied. Laminin-511 was shown to protect islets against cytokine-induced death by interacting with β1 integrins and activating pro-survival Akt signaling. Pro-survival signaling was mediated by reduced activity of protein kinase Cδ (PKCδ), a key mediator of cytokine-induced β-cell death, at the cell membrane in the presence of laminin via reduced levels of diacylglycerol (DAG), a canonical activator of PKCδ. Taken together, these results demonstrate that laminin-511 is an essential factor in protecting β-cells against cytokine-induced death by downregulation of membrane DAG, inhibiting activation of pro-apoptotic PKCδ. Our results suggest that loss of ECM in T1D may make β-cells more susceptible to cytokine-induced death by increasing activation of PKCδ.

**Highlights:**

- Laminin-511 improves islet survival under cytokine treatment
- Pro-survival Akt is upregulated by laminin-511
- Pro-apoptotic PKCδ activity is downregulated at the cell membrane by laminin-511
- Laminin-511 decreases membrane DAG levels leading to reduced PKCδ activation
- Laminin-511 is an essential ECM component for islet survival during type 1 diabetes

## Introduction

The pancreatic islet of Langerhans is a micro-organ composed of endocrine cells that regulate blood glucose and other nutrients through hormone secretion. Among these, β cells are the primary cell type that produce insulin in response to elevated blood glucose. The islet is surrounded by a specialized extracellular matrix (ECM), a complex network of diverse molecules that transmit biochemical signals essential for islet viability and function [1–5]. Overall, the pancreas ECM can be classified into two main components: the peri-islet basement membrane (BM) and the interstitial matrix (IM) [5]. The BM is a thin layer that encapsulates the islet, and the IM makes up the bulk of the pancreas ECM between the islets [5]. The BM is composed of collagen IV, laminin, and heparan sulfate proteoglycans (HSPGs) [2,3,5]. These molecules interact with transmembrane integrin receptors and guide many cellular processes, including cell survival, proliferation, and in the β-cell insulin secretion [4,5]. Laminin is an important component of the peri-islet BM that binds to β1-integrin receptors to enhance islet survival and protect against apoptosis [1,4,6,7]. For example, research with primary islets from rodents has shown that culturing on laminin promotes increased survival and proliferation and protects against cytokine-induced apoptosis via interactions with cell-surface integrins [4,7–9]. The laminin-511 isoform found surrounding human islets is composed of the α5, β1, and γ1 chains and can bind α3β1, α6β1 and α6β4 integrins to activate downstream molecules like focal adhesion kinase (FAK) [4,6,10]. FAK activation triggers the phosphatidylinositol (PI) 3-kinase/protein kinase B (PI3K/Akt) pathway that is known to enhance cell survival [11–14]. While it has been shown in several *in vitro* experiments that laminin enhances β-cell and isolated islet survival, the exact mechanisms behind these signaling pathways are not well understood.

Although type 1 diabetes (T1D) accounts for only 5-10% of all diabetes cases, its global incidence is projected to increase nearly 61% by 2040 [15]. Currently, individuals with T1D rely on continuous glucose monitoring and exogenous insulin therapy to regulate blood glucose levels. However, this approach fails to replicate normal insulin dynamics and is often initiated only after substantial β-cell loss [16]. In T1D, loss of the peri-islet ECM has been shown to occur, even prior to diagnosis before overt symptoms occur [2,5]. As T1D progresses, activated T cells and macrophages secrete proteases that degrade pancreatic ECM molecules like laminin [2]. At the same time, infiltrating T-lymphocytes and macrophages produce elevated levels of pro-inflammatory cytokines, including tumor necrosis factor-α (TNF-α), interleukin-1β (IL-1β), and interferon γ (IFN-γ), that work synergistically to induce β-cell stress, dysfunction, and death as well as production of additional ECM degrading proteases, including matrix metalloproteinase 3 (MMP3) [17–20]. Cytokine-induced β-cell apoptosis is mediated by multiple intracellular signaling pathways, including activation of pro-apoptotic protein kinase C δ (PKCδ) through phosphorylation of its Tyr^311^ domain [21–23]. PKCδ, a member of the novel PKC family, is activated in part by diacylglycerol (DAG), a lipid second messenger that binds to the C1 domain of immature PKCδ, facilitating its recruitment to the plasma membrane for subsequent phosphorylation and activation [22,24,25]. Our previous studies have shown that activated PKCδ drives cytokine-induced β-cell death by nuclear translocation and activation of downstream mediators of apoptosis like c-Jun N-terminal kinase (JNK) and Bax [23]. Therefore, T1D progression involves both activation of cytokine-induced PKCδ signaling and loss of laminin-mediated ECM signaling. While laminin receptor activity has been linked to PKCδ activity in several cell types, its role in regulating cytokine-induced PKCδ activity β-cells remains unknown [26–28].

Thus, the goal of this study is to investigate how β-cell interactions with laminin regulate islet survival under cytokine-induced stress associated with T1D. To achieve this, we used a biomimetic reverse thermal gel (RTG) encapsulation system that can be functionalized with laminin-511 in order to study its effect on islet survival [29–31]. We then exposed encapsulated MIN6 cells, mouse and human islets to a pro-inflammatory cytokine cocktail consisting of tumor necrosis factor α (TNF-α), interleukin-1β (IL-1β), and interferon γ (IFN-γ) and assessed cell viability and pro-survival signaling mediated by laminin activation of cell-surface integrins. Loss of ECM components early in T1D may contribute to disease pathogenesis by disrupting pro-survival signaling and enhancing β-cell susceptibility to immune-mediated death. Therefore, developing therapies that can intervene prior to extensive β-cell loss requires a deeper understanding of the early signaling events driven by ECM loss that lead to β-cell dysfunction and death. The results from this work will improve our understanding of how ECM remodeling contributes to β-cell death and early T1D progression.

## Results

### RTG synthesis and characterization

To study islet interactions with laminin in a 3D environment, we synthesized a reverse thermal gel (RTG) scaffold that has previously been used for 3D culture of various cell types and can be functionalized with ECM molecules, including laminin, to study their role in guiding tissue function [30,31]. The formation and structure of the PSHU-PNIPAAm (RTG) copolymer were evaluated by ^1^H NMR spectroscopy. Characteristic PNIPAAm peaks were observed at 1.11 and 1.60 ppm, consistent with previously reported PSHU-PNIPAAm spectra and supporting successful grafting of PNIPAAm onto the PSHU backbone (Figure 1A) [29,31]. Additional peaks corresponding to PNIPAAm methine protons and methylene protons adjacent to sulfur- and carbonyl-containing groups were also observed.

**Figure 1:**
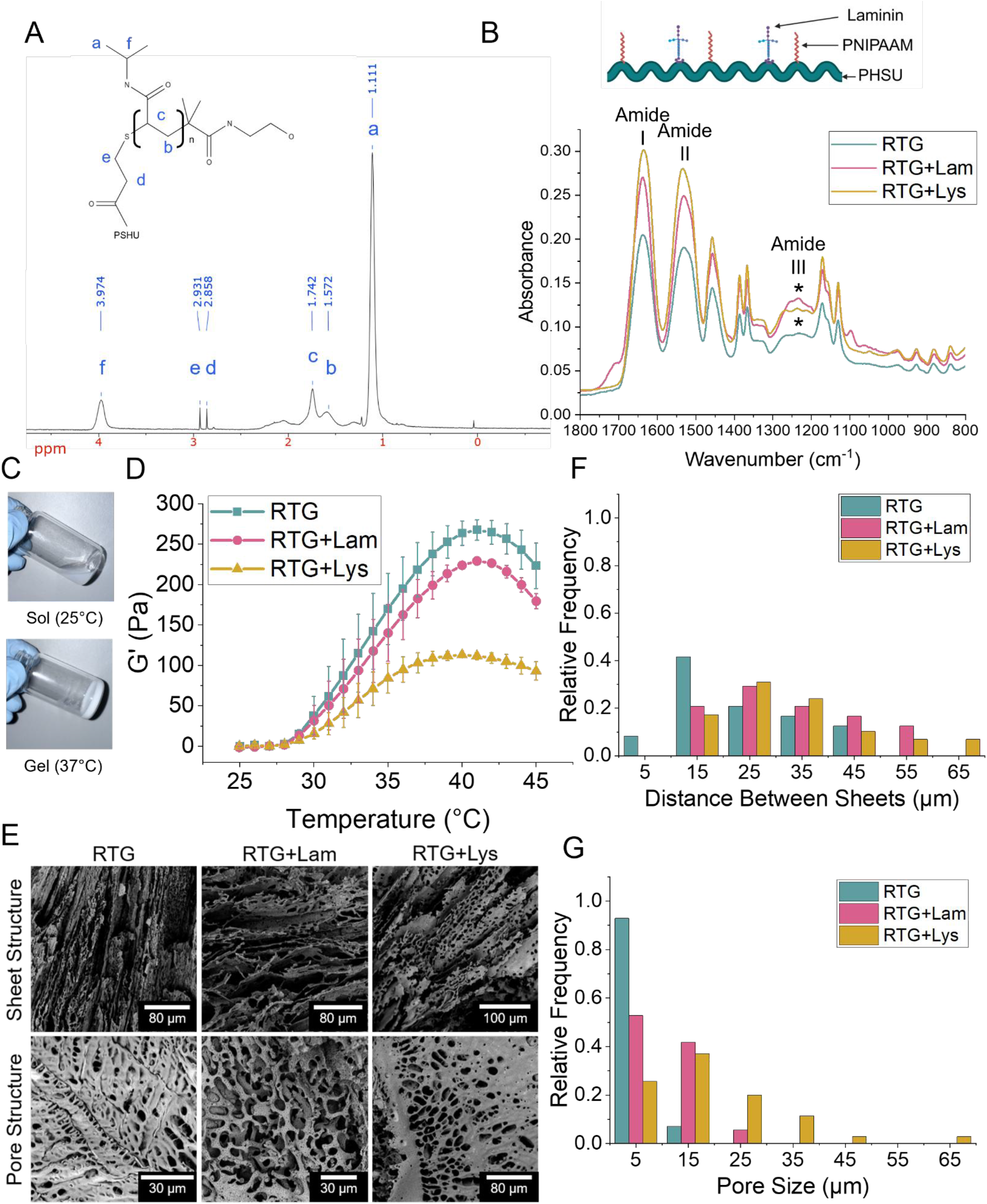
Chemical Characterization of RTG systems. (A) 1H NMR spectrum of PSHU-PNIPAAm. Characteristic PNIPAAm peaks are observed at 1.11 and 1.60 ppm, corresponding to (a) methyl and (b) methylene protons, respectively. Additional peaks (c) and (f) at 1.74 and 3.97 ppm are consistent with the PNIPAAm backbone methine proton and the isopropyl methine proton, respectively. Peaks (d) and (e) at 2.86 and 2.93 ppm are consistent with methylene protons adjacent to sulfur- and carbonyl-containing groups. (B) FTIR spectra of RTG, RTG+Lam, and RTG+Lys. RTG+Lam and RTG+Lys show increased amide I, II and III band compared to RTG, including amide III vibrations (*) at 1230-1280 cm-1, consistent with laminin and lysine functionalization. Top panel created in BioRender. (C) Representative images depicting the 5 wt% RTG reversible phase transition from 25°C to 37°C. (D) Temperature-dependent elastic modulus of 5 wt% RTG, RTG+Lam, and RTG+Lys. (E) SEM images of the dual structure of RTG and RTG+Lam. (F) Relative frequency of distance between sheets and (G) pore sizes of RTG and RTG+Lam samples as measured from the images represented in E. Error bars represent the mean±SEM (n=3).

Laminin-511 was obtained from purified HEK-293 conditioned media by column chromatography and western blot analysis confirmed its presence in the purified sample (Supplemental Figure 1A). The RTG was functionalized with laminin-511 and poly-L-lysine via EDC/NHS chemistry to yield RTG+Lam and RTG+Lys, respectively. The ^1^H NMR spectrum of RTG-Lam retained the characteristic peaks observed in the RTG (Supplemental Figure 1B). The ^1^H NMR spectrum of RTG-Lys was similar to that of RTG but showed an additional triplet at 3.95 ppm, consistent with the α proton of poly-L-lysine (Supplemental Figure 1C) [30].

To confirm attachment of laminin and lysine to the PSHU backbone as shown in the schematic at the top of Figure 1B, FTIR spectra were collected for RTG+Lam and RTG+Lys compared to RTG alone. The FTIR spectra show increased amide I (C=O stretching), amide II (N-H bending and C-N stretching), and amide III (C-N stretching and N-H bending) bands compared to unmodified RTG (Figure 1B) confirming the addition of laminin-511 and lysine respectively [30]. Amide III vibrations were observed at 1220-1260 cm^-1^ in RTG-Lam and RTG-Lys, consistent with previously published spectra for RTG-Lam, and RTG-Lys [30]. Further, previously published work by Peña et al. has demonstrated via thermogravimetric analysis that both the laminin and lysis are functionally attached to the PSHU backbone rather than simply mixed into the RTG solution [30]. The 5 wt% RTG system underwent a sol-to-gel phase transition as temperature increased, whereby the polymer solution turned into a physical gel (Figure 1C). During this transition, the elastic modulus increased sharply between 25 and 33°C, consistent with previously reported thermal gelation behavior of RTG (Figure 1D) [30,31]. Additionally, the coefficient of variance of the storage modulus for the 3 batches of RTG-Lam at 40C was 1.3, suggesting that the degree of functionalization of laminin into the RTG was very consistent from batch to batch.

The surface morphology of 5 wt% RTG, RTG-Lam and RTG-Lys was studied using SEM. Cross-sectional images of the gels revealed highly porous and interconnected sheets (Figure 1E). Quantification from representative SEM images showed a shift toward slightly smaller pore sizes and sheet distances in RTG, while RTG-Lam and RTG-Lys showed distributions shifted towards slightly larger pore sizes and sheet distances (Figure 1F and G) consistent with the addition of bulky laminin and lysine to the RTG backbone [30].

Collectively, characterization of the RTG systems supports successful formation of the thermosensitive PSHU-PNIPAAm copolymer and functionalization of the scaffold with laminin-511 and poly-L-lysine for the study of islet interactions with laminin-511.

### Encapsulated MIN6 cells, mouse islets, and human islets interact with laminin-511 functionalized RTG

To investigate islet interactions with laminin in the RTG+Lam scaffold, we encapsulated isolated mouse islets in 3D scaffolds and visualized the encapsulated islets by immunofluorescence staining of laminin and cell nuclei (Figure 2A). Laminin-511 staining was observed throughout the RTG+Lam scaffold surrounding the islet cells, confirming the presence of laminin within the encapsulation environment (Figure 2A). To determine if β-cells express integrin receptors that can bind to laminin-511 in the RTG environment, we stained for β1 integrins in fixed MIN6 cells, mouse and human islets [10]. In MIN6 cells, β1 integrin was localized to the cell borders, particularly at sites of cell-cell contact and along the bottom surface of cells in contact with the culture dish (Figure 2B). We then confirmed integrin staining in isolated mouse islets, the primary model used for subsequent functional studies. While β1 integrin remained enriched at the mouse islet periphery and at cell-cell interfaces, it appeared less abundant and widespread compared to MIN6 cells (Figure 2C). To evaluate whether this expression pattern was preserved in β-cells within the native pancreatic islet environment, we performed immunohistochemistry staining of a C57Bl/6 mouse pancreas (Supplemental Figure 2). Our results revealed β1 integrin localization around islets and adjacent to insulin- and glucagon-positive cells, indicating that integrins are present in mouse β-cells *in situ* with similar expression patterns in isolated islets. In contrast, human islets exhibited stronger β1 integrin staining than either MIN6 cells or mouse islets, which was primarily located at the islet periphery and surrounding individual endocrine cells, including both β and α cells (Figure 2D). This data shows that MIN6 cells, mouse and human islets express β1 integrins that can interact with molecules within their encapsulation environment. Focal adhesion kinase (FAK) is activated by phosphorylation following integrin binding to extracellular ligands and mediates downstream survival signaling [32]. To determine if encapsulated islets can bind to laminin-511 in the RTG scaffold, we encapsulated MIN6 cells, mouse and human islets in RTG, RTG+Lam and RTG+Lys for 24 h and analyzed FAK phosphorylation at Tyr^397^ by western blot quantified as p-FAK/FAK and normalized to the RTG only samples for each experimental replicate (Figure 3A-F). MIN6 cells encapsulated in RTG+Lam and RTG+Lys had increased FAK activity, as measured by normalized p-FAK/FAK, compared to those in RTG (Figure 3A and B). Similarly, mouse islets in RTG+Lam and RTG+Lys had increased FAK activity compared to islets in RTG alone (Figure 3C and D). Human islets encapsulated in RTG+Lam had significantly higher FAK activity than in RTG (Figure 3E and F). Islets encapsulated in RTG+Lys also showed a modest increase in FAK activity relative to RTG. As a positive control of FAK activation upon β1 integrin binding, mouse islets were treated with the β1 integrin agonist pyrintegrin for 24 h and we observed increased FAK activity compared to untreated islets (p=0.149, Supplemental Figure 3A and B). As a negative control, human islets were treated with the anti-β1 integrin function-blocking antibody, AIIB2, 30 minutes before encapsulation in RTG+Lam and showed significantly lower FAK activity compared to islets in RTG+Lam (Figure 3E and F, p=0.027), supporting a role for the β1 integrin in directly binding to laminin-511 and activating intracellular signaling cascades. In summary, islets encapsulated in RTG functionalized with laminin-511 can interact with laminin in the scaffold to activate intracellular signaling cascades, including FAK.

**Figure 2:**
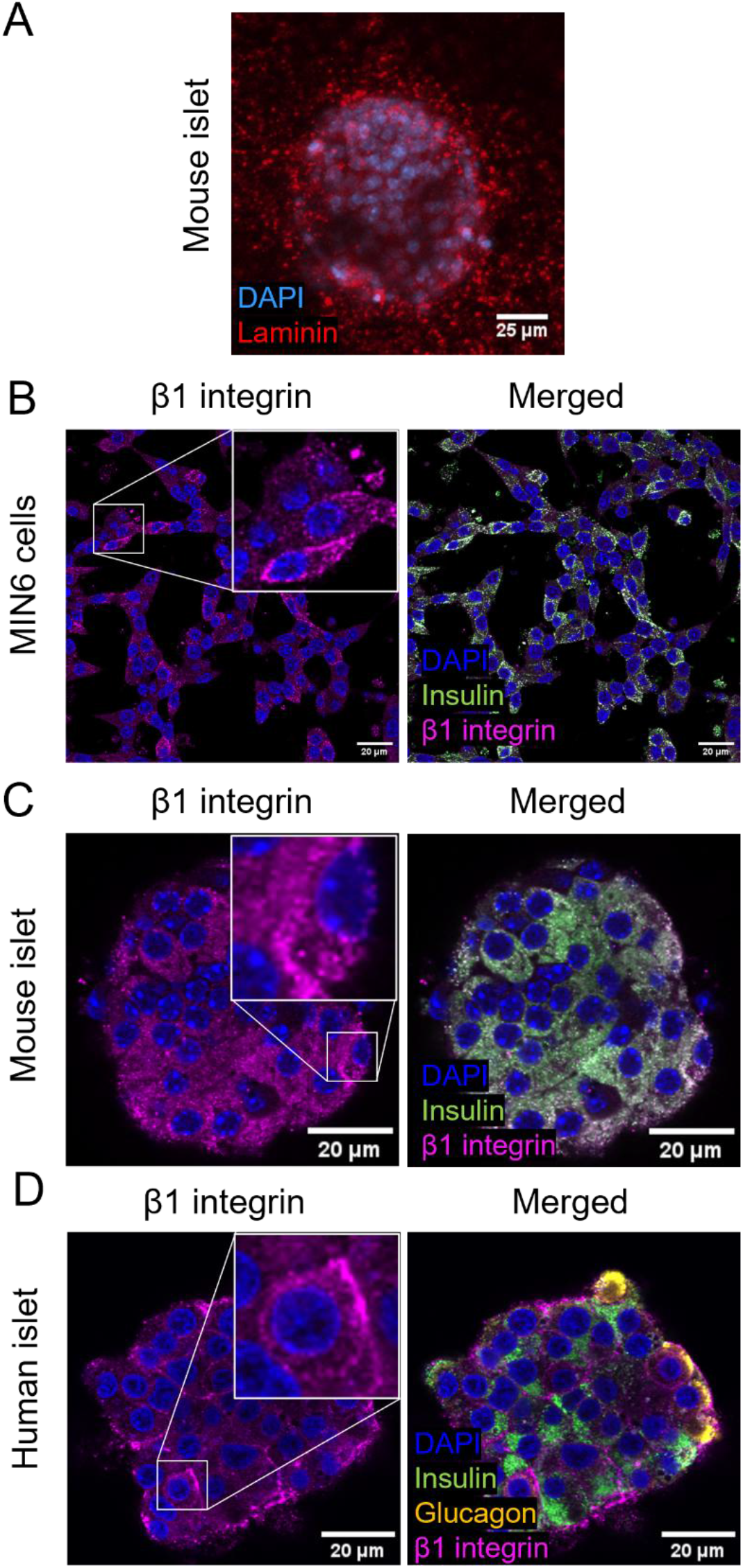
Distribution of β1 integrins around islet cells. (A) IHC image of a mouse islet encapsulated in RTG-LAM stained with DAPI (blue) and laminin (red). IHC images of fixed MIN6 cells at the basal focal plane (B), a fixed B6 mouse islet (C), and a fixed human islet (D) stained with DAPI (blue), insulin (green), glucagon (yellow) and β1 integrin (magenta).

**Figure 3:**
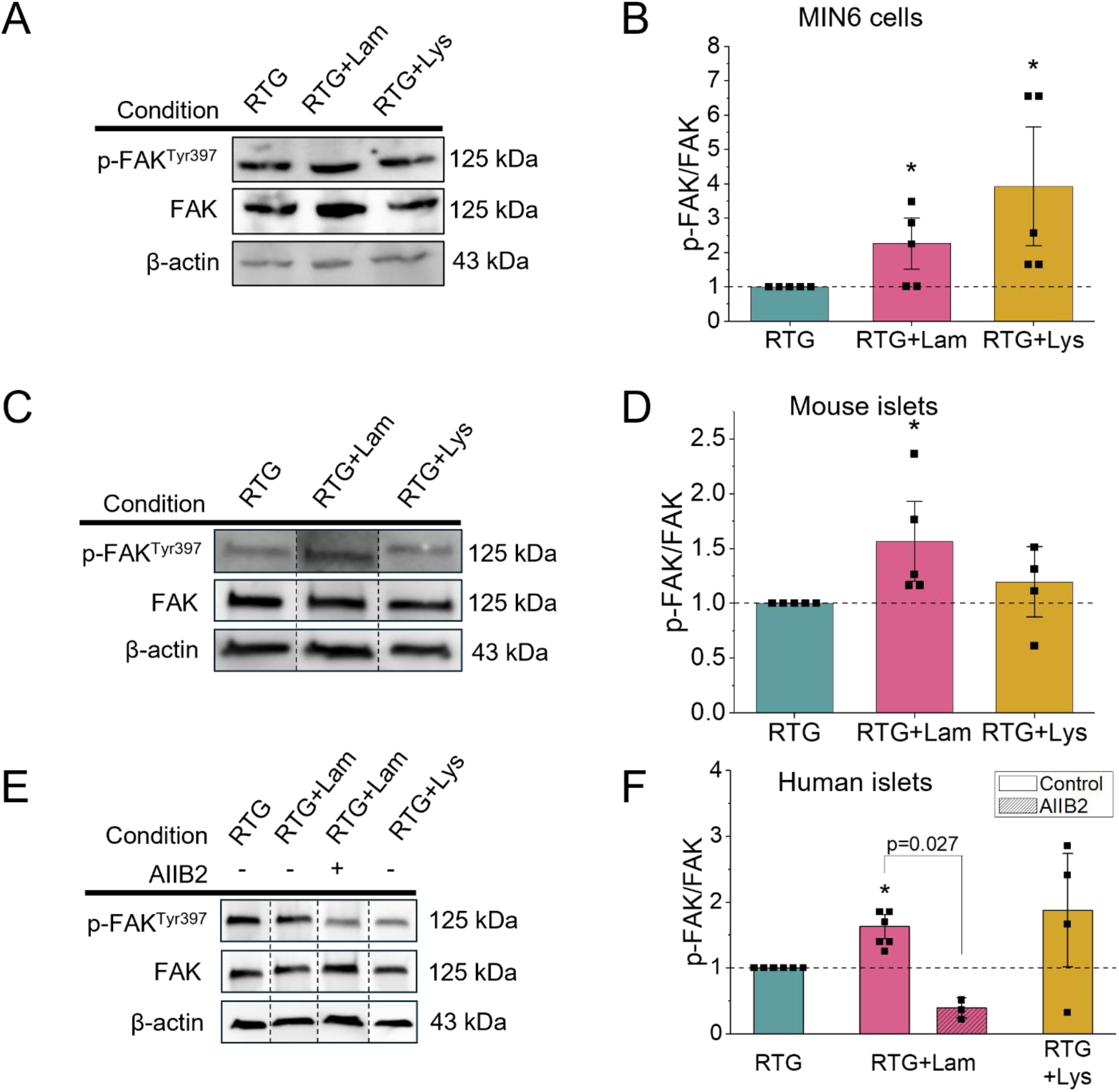
Islet cells interact with the RTG matrix and laminin. (A) Representative western blot images of p-FAK, FAK and β actin present in MIN6 cells encapsulated in RTG, RTG+Lam, or RTG+Lys for 24 h. (B) Western blot quantification of p-FAK/FAK in MIN6 cells encapsulated in RTG, RTG+Lam or RTG+Lys, with data normalized to RTG (n=5). (C) Representative western blot images of p-FAK, FAK and β actin present in B6 mouse islets encapsulated in RTG, RTG+Lam, or RTG+Lys for 24 h. Blot images were cropped for presentation. The vertical dashed line indicates nonadjacent lanes from the same blot and exposure. (D) Western blot quantification for p-FAK/FAK in mouse islets encapsulated in RTG (n=5), RTG+Lam (n=5) or RTG+Lys (n=4), with data normalized to RTG. (E) Representative western blot images of p-FAK, FAK and β actin present in human islets encapsulated in RTG, RTG+Lam, or RTG+Lys for 24 h and either untreated or treated with the anti-β1 integrin antibody AIIB2. Blot images were cropped for presentation. The vertical dashed line indicates nonadjacent lanes from the same blot and exposure. (F) Western blot quantification for p-FAK/FAK in human islets encapsulated in RTG (n=6), RTG+Lam untreated (n=6), RTG+Lam + AIIB2 (n=3) or RTG+Lys (n=4), with data normalized to RTG. Error bars represent the mean±SEM. * Indicates a significant difference using a one-sample t-test against a theoretical mean of 1. p-value <0.05 is significant as determined by ANOVA.

### Laminin-511 protects against cytokine-mediated islet death

To determine if islet interactions with laminin-511 protect against cytokine-induced death, we assessed islet viability within the RTG scaffold either alone, with laminin-511, or with lysine as a control for non-specific integrin activation. Human islets encapsulated in RTG, RTG+Lam, and RTG+Lys, including islets encapsulated in RTG-Lam had no significant differences in cell death compared to unencapsulated islets (Figure 5 A and B). After treatment with a cocktail (1x) of human recombinant cytokines composed of 10 ng/ml TNF-α, 5 ng/ml IL-1β, and 100 ng/ml IFN-γ for 24 h, cell death significantly increased for islets encapsulated in RTG, RTG+Lys, and unencapsulated islets (Figure 5 A and B). Human islets encapsulated in RTG+Lam and treated with cytokines showed significantly less cell death relative to islets in RTG alone (p=0.019, Figure 4B and C). Additionally, islets encapsulated in RTG+Lam and treated with the β1 integrin inhibitor AIIB2 were not protected against cytokine-induced death and showed similar cell death as islets encapsulated in RTG alone (Figure 4C). Similar to human islets, mouse islets that were encapsulated in RTG, RTG+Lam, and RTG+Lys for 24 hours showed no significant differences in cell death compared to unencapsulated islets (Supplemental Figure 4, Figure 4D and E). Treatment with a cocktail (0.1x) of mouse recombinant cytokines composed of 1 ng/ml TNF-α, 0.5 ng/ml IL-1β, and 10 ng/ml IFN-γ for 24 h significantly increased cell death for unencapsulated islets and islets in RTG alone and RTG+Lys. However, mouse islets that were encapsulated in RTG+Lam had similar viability as untreated controls. Additionally, islets in RTG+Lam showed significantly less cell death with cytokine treatment compared to islets encapsulated in RTG alone (p=0.049) or compared to RTG+Lys (Supplemental Figure 4, Figure 4D and E). Taken together, this data supports that laminin interacts with islets via the β1 integrin to protect against pro-inflammatory cytokine induced cell death.

**Figure 4:**
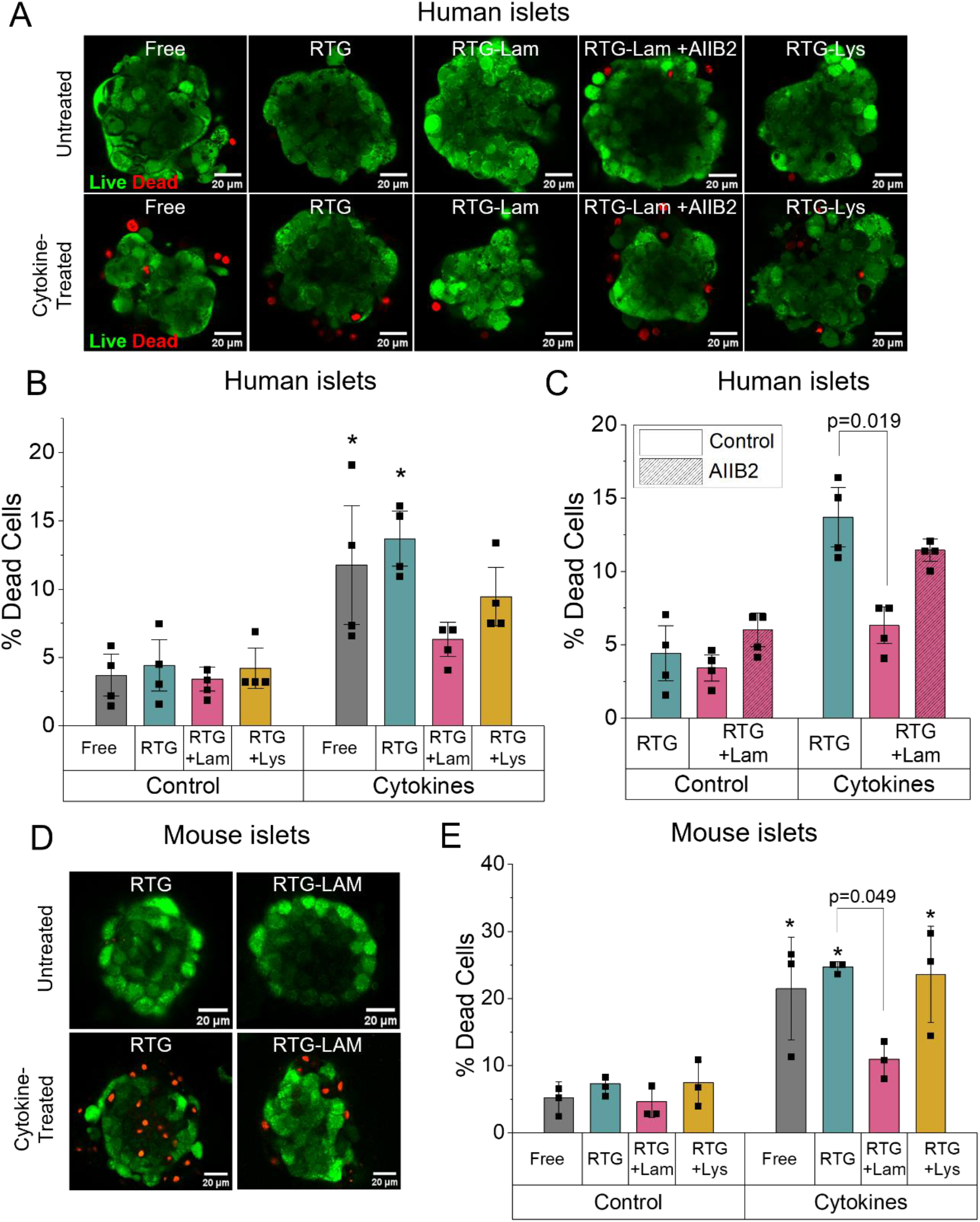
Laminin improves human islet viability under cytokine stress. (A) Representative images of human islet viability in no RTG and 5 wt% RTG, RTG+Lam, and RTG+Lys either untreated or treated with 1X cytokines 24 h or the anti-β1 integrin antibody AIIB2 for 30 min before encapsulation. (B) Percent dead cells in control or cytokine-treated human islets either freely cultured or encapsulated in 5 wt% RTG, RTG+Lam, or RTG+Lys for 24 h (n=4). (C) Percent dead cells in control, cytokine-treated or AIIB2-treated human islets either freely cultured or encapsulated in 5 wt% RTG, RTG+Lam, or RTG+Lys for 24 h (n=4). (D) Representative images of mouse islet viability in 5 wt% RTG and RTG+Lam either untreated or treated with cytokines for 24 h. (D) Percent dead cells in control, 0.1X cytokine-treated mouse islets either freely cultured or encapsulated in 5 wt% RTG, RTG+Lam, or RTG+Lys for 24 h (n=3). Data represents the mean±SEM. * Indicates a significant difference from the respective untreated islets and p-value <0.05 is significant as determined by ANOVA.

**Figure 5:**
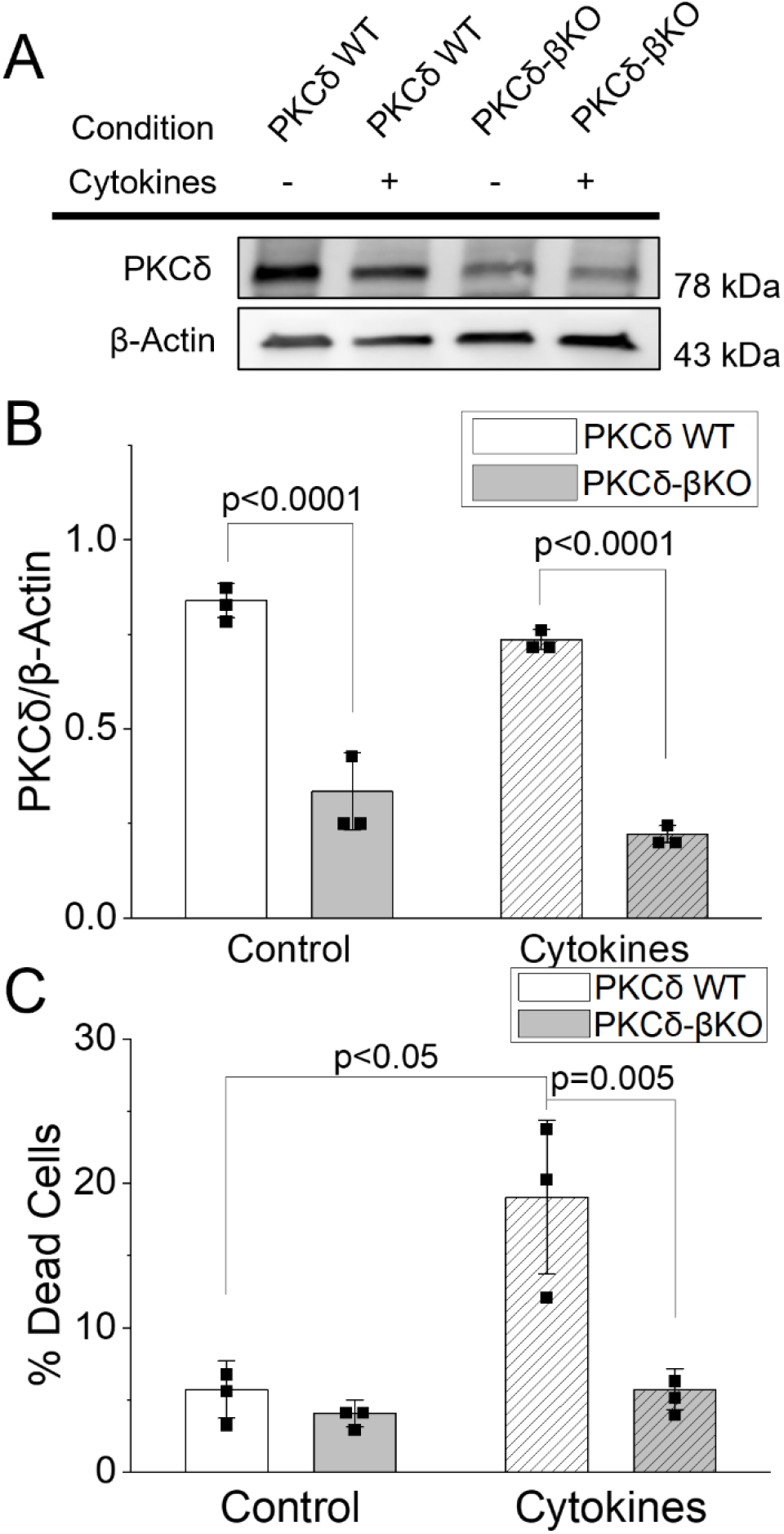
PKCδ knockout protects against cytokine-induced death. (A) Representative western blot images of PKCδ and β actin present in PKCδWT and PKCδKO mouse islets either untreated or treated with cytokines for 24 h. (B) Western blot quantification for PKCδ normalized to β-actin in PKCδWT and PKCδKO mouse islets (n=3). (C) Percent dead cells in PKCδWT and PKCδKO mouse islets either untreated or treated with cytokines (n=3). Data represents the mean±SEM. p-value <0.05 is significant as determined by ANOVA.

To further assess the role of decreases in pro-apoptotic PKCδ signaling in mediating cytokine-induced β-cell death, we developed a β-cell specific knockout (KO) of PKCδ on the MIP Cre ER background (PKCδ-βKO) [23]. Isolated PKCδ-βKO and Cre expressing PKCδ WT islets were cultured with or without 1X mouse cytokines for 24 h. PKCδ-βKO islets had significantly less PKCδ protein compared to the WT control in both the untreated and cytokine-treated groups, confirming successful knockout of PKCδ (p<0.001, Figure 5A and B). Upon cytokine treatment, PKCδ-βKO islets had significantly fewer dead cells compared to PKCδ WT islets (p=0.005, Figure 5C). This supports a role for PKCδ in mediating cytokine-induced death.

### Laminin-511 downregulates PKCδ activity at the cell membrane

To further determine if PKCδ plays a role in the mechanism by which laminin-511 protects against cytokine-induced cell death, we employed a membrane-targeted δ-selective PKC activity reporter (δCKAR) [33]. δCKAR is a FRET-based biosensor that we transfected into MIN6 cells to measure PKCδ activity specifically at the plasma membrane as shown in Figure 6A with YFP and CFP expression localized to the cell membrane. Since increased δCKAR activity corresponds to a decrease in the FRET/CFP ratio, increased PKCδ activity was reported as the inverse FRET ratio such that increased ratio values represent increased PKCδ activity as shown in representative ratiometric images of the FRET sensor in Figure 6B and quantification of inverse FRET efficiency in Figures 6C and D. During the first 5 minutes of imaging, the cells were left untreated and there was no observed change in PKCδ activity (represented as normalized 1/E, Figure 6B and C). After 5 minutes, the cells were treated with 1X mouse cytokine cocktail, and PKCδ activity increased in MIN6 cells encapsulated in RTG. However, there was a significant decrease in PKCδ activity in MIN6 cells encapsulated in RTG+Lam compared to those encapsulated in RTG alone after cytokine treatment (Figure 6B, C, and D, p=0.036). Treatment with the pan PKC activator phorbol 12-myristate 13-acetate (PMA) at 30 minutes increased the activity of PKCδ in MIN6 cells encapsulated in RTG, which remained higher than that of cells encapsulated in RTG+Lam (Figure 6B, C, and D, p=0.023). Overall, these results suggest that laminin interactions with islet β-cells downregulates PKCδ activity at the MIN6 cell membrane.

**Figure 6:**
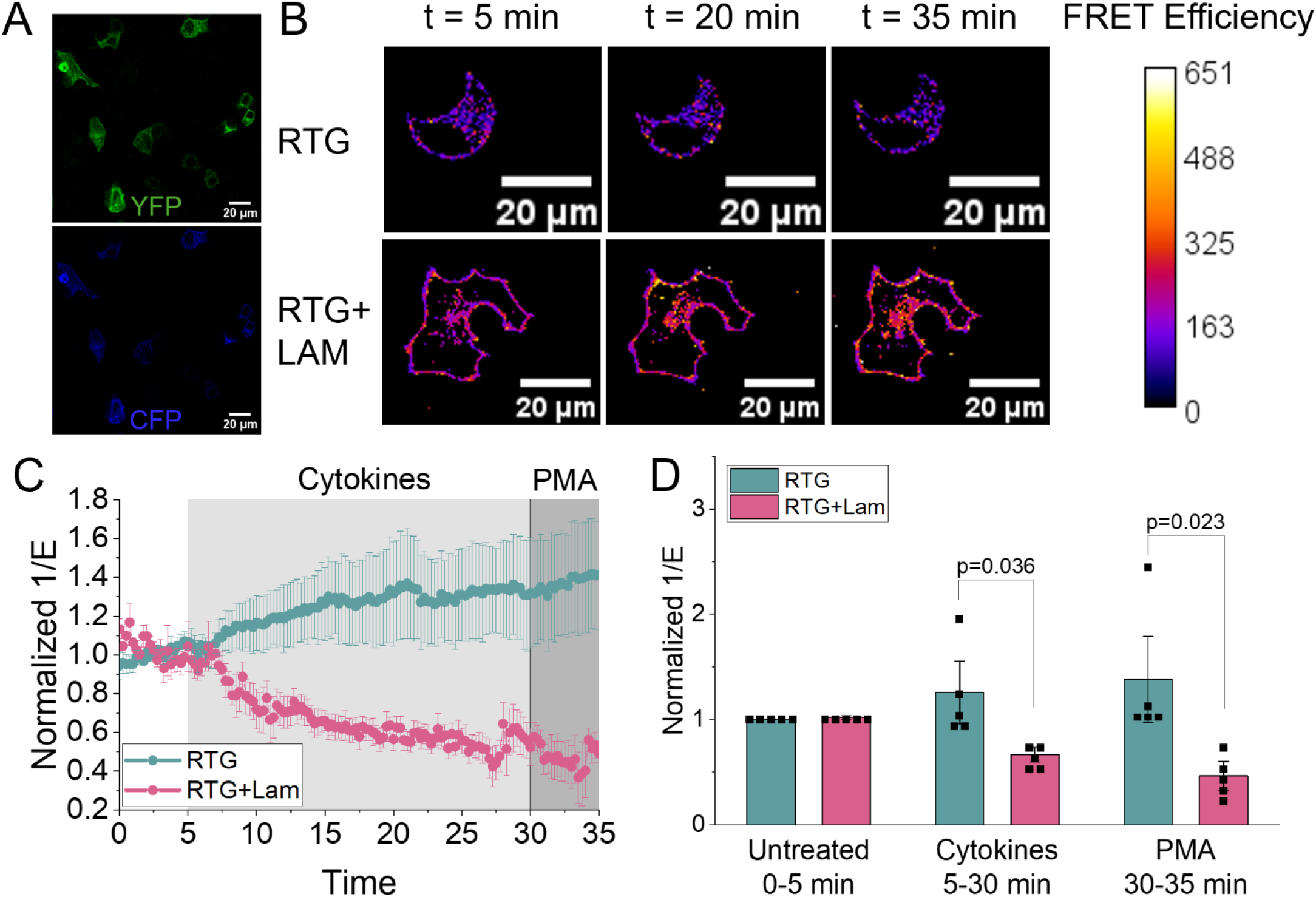
Laminin attenuates PKCδ membrane FRET activity. (A) Representative images of YFP (top, yellow) and CFP (bottom, blue) expression in MIN6 cells after transfection with the δCKAR PKCδ activity biosensor. (B) Representative FRET/CFP ratio images of MIN6 cells encapsulated in RTG and RTG+Lam. FRET efficiency is displayed as a pseudocolor LUT, where darker colors indicate lower FRET/CFP ratios and brighter colors indicate higher FRET/CFP ratios. (B) Average inverse FRET efficiency (1/E) of a membrane-targeted PKCδ activity biosensor in MIN6 cells encapsulated in RTG and RTG+LAM and treated with 1X cytokines and 300 μM of the PKCδ activator PMA as indicated (n=5). Data are normalized to 1/E from 2-5 minutes prior to treatments. (C) Inverse FRET efficiency (1/E) averaged over the indicated time intervals normalized as in A. Error bars represent the mean±SEM and p-value <0.05 is significant as determined by ANOVA.

### Laminin-511 activates pro-survival Akt in MIN6 cells and human islets mediated by PKCδ activity

The PI3K/Akt pathway is known to play an important role in regulating β-cell survival [13,34]. Once activated, Akt, the key downstream effector of the PI3K pathway, can directly inhibit various apoptotic stimuli [35]. To determine if pro-survival Akt signaling is upregulated with laminin-511 and β1 integrin activation, we measured levels of phosphorylated Akt at Thr^308^ normalized to Akt as a measure of Akt activity via western blot. In mouse islets treated with the β1 integrin agonist pyrintegrin for 24 h we saw an increase in Akt activity compared to untreated islets (Supplemental Figure 5A and B). Further, to determine whether laminin-511 induces activation of Akt in the β-cell, we encapsulated MIN6 cells, mouse and human islets in RTG, RTG+Lam and RTG+Lys and treated them with 0.1 mouse or 1X human cytokines for 24 h (Figure 7A-D). For MIN6 cells encapsulated in RTG, RTG+Lam, or RTG+Lys and left untreated for 24 h, Akt activity as measured by p-Akt/Akt was increased in RTG-Lam compared to RTG (Figure 7A and B). This increase was maintained during the 1 h cytokine treatment (Supplemental Figure 5C and D) as well as the 24 h cytokine treatment (Figure 7A and B). In contrast MIN6 cells in RTG+Lys and RTG alone showed decreased Akt activity with cytokine treatment (Figure 7A and B). Similar to MIN6 cells, untreated human islets in RTG+Lam showed significantly increased Akt activity compared to those in RTG or RTG+Lys (Figure 7C and D). Additionally, cytokine-treated human islets in RTG+Lam also showed increased Akt activity compared to those in RTG and RTG-Lys. In contrast, both untreated and cytokine-treated mouse islets encapsulated in RTG+Lam did not show a significant difference Akt activity compared to RTG and RTG+Lys conditions (Supplemental Figure 5E and F). Finally, human islets were treated with the anti-β1 integrin function-blocking antibody AIIB2 30 min before through 24h after encapsulation in RTG+Lam and showed lower Akt activity compared to islets in RTG+Lam without AIIB2 treatment (p=0.049, Figure 5C and D). Altogether, this data affirms laminin’s ability to activate the pro-survival pathway involving Akt in β-cells via activation of the β1 integrin.

**Figure 7:**
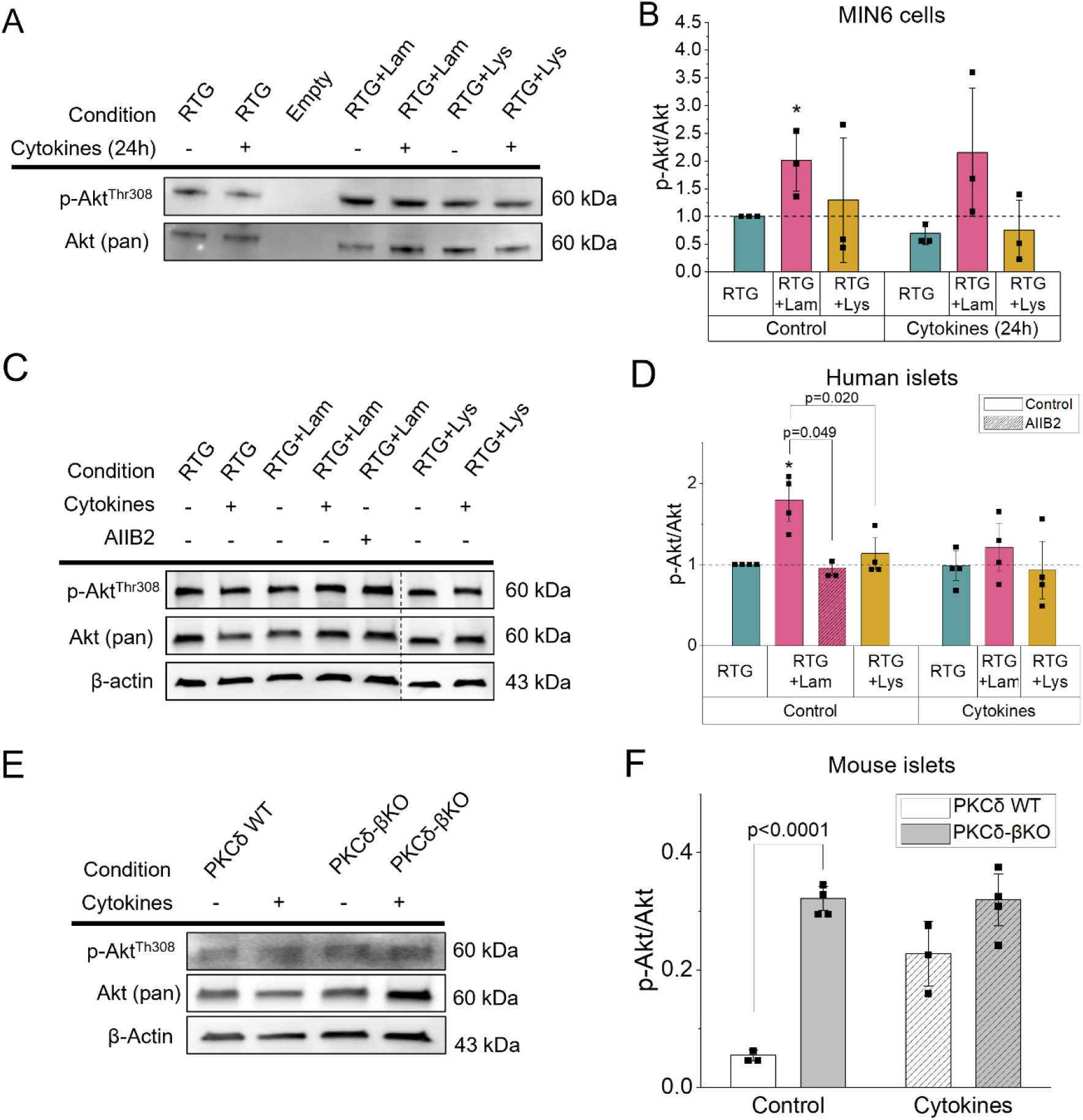
Laminin promotes Akt signaling. (A) Representative western blot images of p-Akt and Akt present in MIN6 cells encapsulated in RTG, RTG+Lam, or RTG+Lys and either untreated or treated with a cytokine cocktail for 24 h. (B) Western blot quantification for p-Akt/Akt in MIN6 cells (n=3). Data is normalized to RTG (control). (C) Representative western blot images of p-Akt, Akt and β actin present in human islets encapsulated in RTG, RTG+Lam, or RTG+Lys and either untreated or treated with a cytokine cocktail for 24 h or the anti-β1 integrin antibody AIIB2 for 30 min before encapsulation. Blot images were cropped for presentation. The vertical dashed line indicates nonadjacent lanes from the same blot and exposure. (D) Western blot quantification for p-Akt/Akt in human islets encapsulated in RTG untreated (n=4), RTG+Lam untreated (n=4), RTG+Lam + AIIB2 (n=3), RTG+Lys untreated (n=4), RTG + cytokines (n=4), RTG+Lam + cytokines (n=4), or RTG+Lys + cytokines (n=4). (E) Representative western blot images of p-Akt, Akt, and β actin present in PKCδWT and PKCδKO mouse islets either untreated or treated with cytokines for 24 h. (F) Western blot quantification for p-Akt/Akt in PKCδWT islets untreated (n=3), PKCδKO islets untreated (n=4), PKCδWT islets + cytokines (n=3), PKCδKO islets + cytokines (n=4). Data in B and D is normalized to RTG (control). Error bars represent the mean±SEM. * Indicates a significant difference using a one-sample t-test against a theoretical mean of 1. Differences among treatment groups were assessed separately within the control and cytokine-treated conditions using one-way ANOVA. Statistical significance was defined as p <0.05.

To determine the effect of PKCδ on Akt activation, phosphorylation of Akt at Thr^308^ in isolated PKCδ-βKO and PKCδ WT islets was measured by western blot as above. Untreated PKCδ-βKO islets showed significant upregulation of Akt compared to untreated Cre expressing PKCδ WT islets (p=0.008, Figure 7E and F). While no significant difference was detected between PKCδ-βKO and PKCδ WT islets upon treatment with cytokines (p=0.102), PKCδ-βKO islets had slightly higher levels of Akt activity (Figure 7F). Combined, this data indicates that PKCδ plays an important role in regulating the PI3K/Akt pathway in β-cells and may partly mediate the protective effects of laminin in cytokine treated islets.

**Figure 8:**
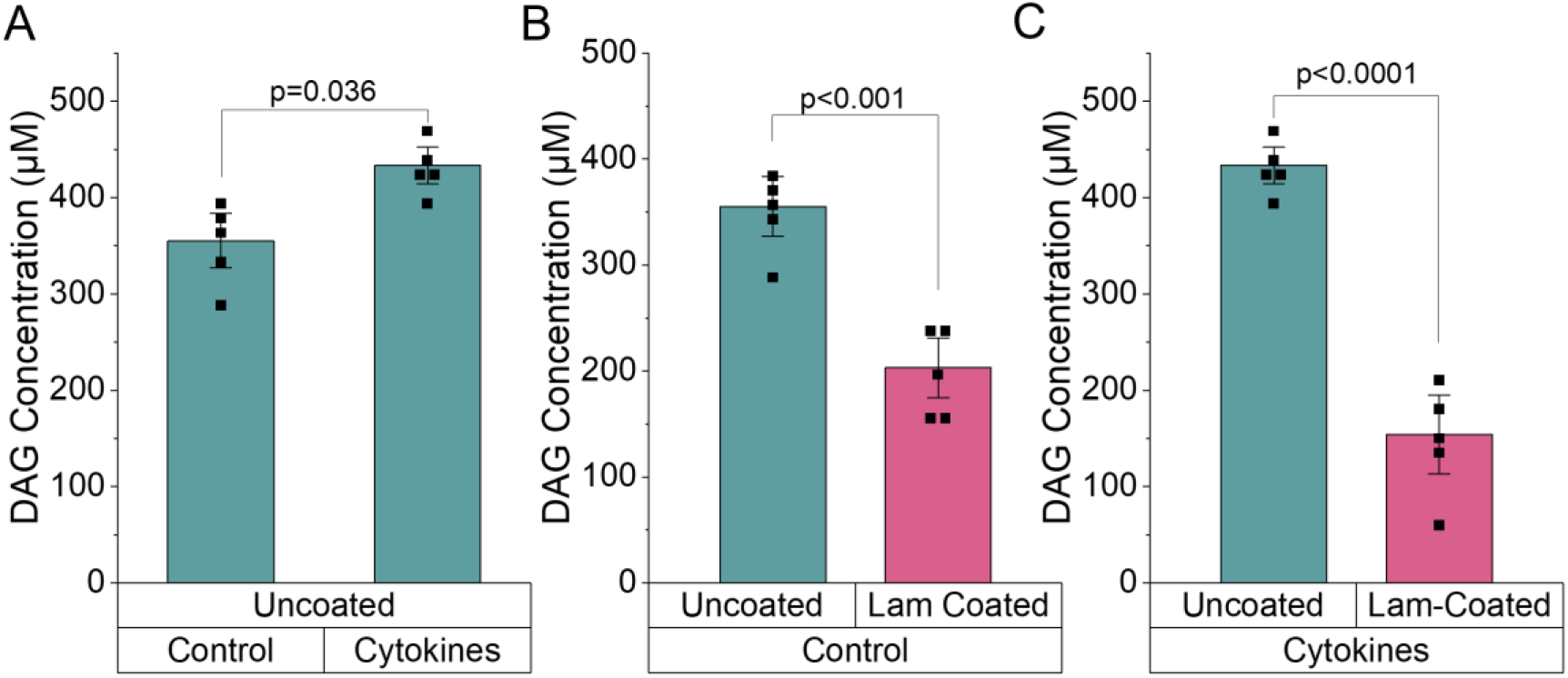
Diminished DAG content with laminin. Amount of DAG present in MIN6 cells grown on (A) tissue culture-treated (uncoated) dishes and treated with or without cytokines, (B) cultured on uncoated or Lam coated dishes, or (C) cultured on uncoated or Lam coated dishes and treated with cytokines for 24 h. Error bars represent the mean±SEM and p-values are significant as determined by student’s T-test in A and ANOVA in B and C.

### Laminin-511 downregulates DAG in MIN6 cells

To investigate the mechanism by which laminin-511 downregulates PKCδ at the cell membrane, we measured levels of DAG, a lipid second messenger that acts as a canonical PKCδ activator, in MIN6 cells cultured with and without laminin-511 and either untreated or treated with 1X mouse cytokines for 24 h. Analysis of DAG levels reveals that cytokines increased the amount of DAG in the cells, consistent with previously published findings (p=0.036, Figure 9A) [36,37]. However, untreated MIN6 cells cultured on laminin-511 had significantly decreased DAG levels compared to cells cultured on standard poly-L-lysine coated tissue culture flasks (p<0.001, Figure 9B). Finally, MIN6 cells cultured on laminin-511 and treated with cytokines had significantly decreased DAG levels compared to cells cultured on standard tissue culture flasks also treated with cytokines (p<0.001, Figure 9C). This result supports the notion that laminin interactions with β-cells alters DAG content and therefore, activation of pro-apoptotic PKCδ signaling under normal and cytokine stressed conditions.

**Figure 9:**
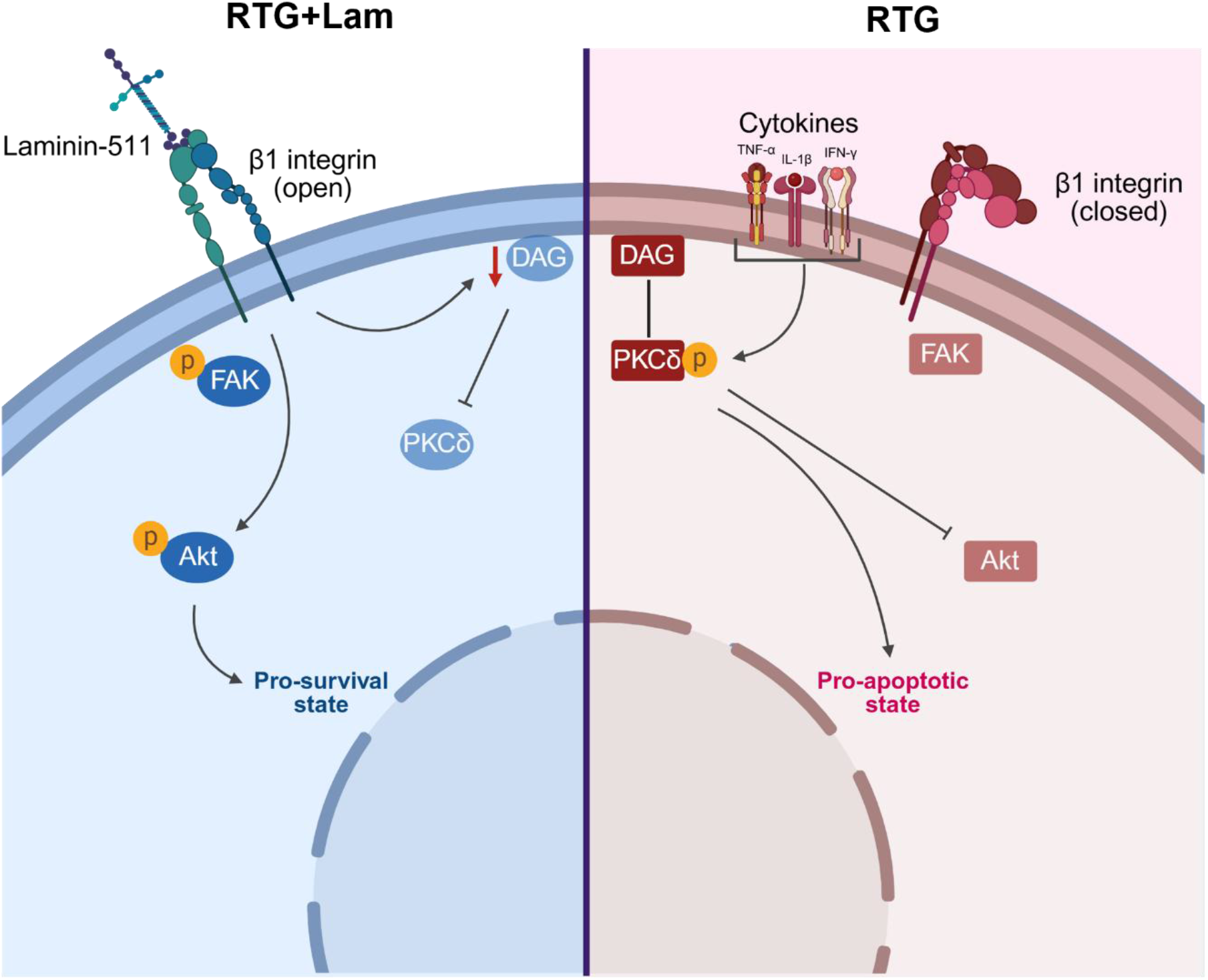
Laminin-511 activates pro-survival signaling pathways within the β cell to protect against cytokines. When present in the RTG scaffold, laminin-511 binds to β1 integrins on the β-cell membrane to activate focal adhesion kinase (FAK) and Akt, promoting a pro-survival state in the cell. Additionally, laminin reduces levels of diacylglycerol (DAG) at the membrane, thereby downregulating pro-apoptotic protein kinase Cδ. When laminin is absent from the RTG scaffold, β1 integrins are typically in the closed conformation and FAK is inactive. DAG binds to the C1 domain of PKCδ to drive its membrane translocation and activation. Treatment with a cytokine cocktail also activates PKCδ, which will downregulate pro-survival Akt and promote a pro-apoptotic state in the cell. Created in BioRender.

## Discussion

Laminin-511 is a peri-islet ECM molecule known to increase pancreatic β-cell survival. In 2D culture, laminin has been shown to activate focal adhesions at the β-cell membrane and upregulate pro-survival Akt; however, the role of loss of laminin signaling and increased β-cell death in T1D has not been explored to date [4,6,7]. Additionally, the signaling cascade between laminin and activation of downstream pro-survival proteins like Akt in the β-cell is not well defined, especially in the context of T1D. The goal of this study was to determine the role of laminin in regulating β-cell survival under conditions associated with T1D and identify the molecular mechanisms behind this protection. We functionalized a biomimetic 3D RTG encapsulation system functionalized with laminin-511 (RTG+Lam) and measured the activity of downstream pro-survival signaling proteins. Our results indicate that laminin activates β1 integrins at the membrane to downregulate pro-apoptotic DAG and PKCδ and upregulate pro-survival Akt to protect against cytokine-induced death in β-cells (Figure 10). As loss of the peri-islet ECM occurs during T1D development, understanding the consequences of loss of laminin-511 on important signaling pathways will provide insight into islet death during T1D.

### RTG is a viable platform for studying laminin interactions with β cells in 3D

For this study, we utilized a 3D biomimetic polymer that could be easily functionalized with ECM molecules and allow for facile extraction of encapsulated islets for downstream analysis making it an ideal platform for studying laminin’s influence on islet apoptosis *in vitro* [29–31].The RTG is composed of interconnected pores and sheets that are much smaller than an islet, providing suitable encapsulation while facilitating the movement of nutrients and waste through the scaffold. The chemical incorporation of laminin and poly-L-lysine increased the pore size and sheet distance, likely due to the bulky size and the repulsion between the hydrophilic groups in both laminin and poly-L-lysine and the hydrophobic groups of the PSHU-PNIPAAm polymer [30]. Islet encapsulation in RTG, RTG+Lam, or RTG+Lys supported islet viability over 24h in culture. Our previous studies using RTG have also shown a high level of diffusivity of both small and large molecules within the scaffold, indicating that RTG encapsulation does not compromise islet integrity or reduce nutrient diffusion [38]. Additionally, we saw similar levels of β-cell death in cytokine treated islets encapsulated in RTG compared to unencapsulated islets, further supporting that the RTG scaffold does not impart any diffusional barriers to oxygen or nutrients. These findings support the use of the RTG system as a platform to study islets in a 3D environment without inducing encapsulation-related damage. To further validate the RTG model for studying islet interactions with laminin, we measured laminin binding to the β1 integrin on the surface of β-cells. Activation of β1 integrins by binding to ECM molecules leads to activation of FAK and other proteins such as extracellular-regulated kinase (ERK), Akt, and the steroid receptor coactivator (Src) family of kinases, all of which are involved in cell survival [3,4,39]. The increase in phosphorylation of FAK in MIN6 cells and islets encapsulated in RTG+Lam affirms that laminin engages and activates β1 integrins which we showed were highly expressed on the surface of the β-cell. This also confirms that covalent attachment of laminin-511 to the RTG backbone did not impact its ability to bind to cell surfaced integrins, making the laminin-511 functionalized RTG an ideal scaffold to investigate ECM interactions with islets in a 3D environment.

### Laminin protects β cells against cytokine-induced death

Similar to previously published results in MIN6 cells and human islets, we found that islet interactions with laminin-511 increase the survival of isolated pancreatic islets treated with an acute high dose of a cytokine cocktail *in vitro* [1,4,6,7]. Additionally, several studies using the function-blocking antibody AIIB2 directed against the β1 integrin subunit have demonstrated decreased cell proliferation and increased apoptosis as a result of perturbed integrin signaling [40,41]. Here we show that treating human islets with AIIB2 prior to RTG+Lam encapsulation reduces FAK activation and increases β-cell death. These results suggest that β1 integrin engagement and activity is critical for maintaining laminin-mediated islet survival, particularly under cytokine stress. Further, we found that activation of pro-survival Akt signaling was dependent on expression levels of β1 integrins across the β-cell and islet models studied. Specifically, human islets expresses the highest levels of β1 integrin on the islet surface and consequently had the largest increase in Akt activity both with and without cytokines compared to islets in RTG alone, while mouse islet has the lowest levels of β1 integrin expression on the cell surface and consequently also had the lowest activity of Akt across all treatments tested. This further supports the role for β1 integrin signaling activated by laminin binding in regulating islet survival. Given that cytokines represent only a subset of the inflammatory milieu associated with T1D, further studies are needed to determine if laminin protects against broader inflammatory stress and autoreactive immune cell mediated β-cell death in T1D. Collectively, our findings support a role for laminin-511 in mediating cytokine-induced β-cell death via interactions with β1 integrins.

### Laminin-511 decreases pro-apoptotic signaling via downregulation of PKCδ activation

In several cell types, laminin receptor activity is linked to PKCδ function; however, the mechanisms regulating PKCδ activity have not been explored [26–28]. PKCδ activation requires translocation of immature and inactive PKCδ to the plasma membrane and binding of DAG to its C1 domain [25,42,43]. Here we have found that PKCδ activity is indirectly regulated by β-cell interactions with laminin mediated through decreased membrane DAG content, as shown in Figure 10. One possible mechanism to explain how laminin binding to β1 integrins regulates DAG content involves diacylglycerol kinase (DGK), which transforms DAG into phosphatidic acid (PA) and thus limits DAG-regulated functions like PKCδ activation [44]. Since integrin signaling has been found to regulate DGK, we propose that laminin binding to integrins on the β cell leads to DGK upregulation and a subsequent decline in DAG [45]. on the reduced DAG content in cells cultured with laminin led to reduced PKCδ activity at the cell membrane, as a reduction in DAG availability has been shown to inhibit PKCδ activation, and this resulted in increased pro-survival Akt signaling and reduced cytokine induced β-cell death [46,47]. While our results demonstrate a decrease in DAG content in cells grown on laminin-coated dishes, one limitation of this platform is that it does not fully recapitulate the 3D microenvironment of intact islets. Therefore, further studies that investigate laminin signaling with DGK and DAG in living pancreas slices from human donors could provide better insight into laminin regulation of PKCδ and broader mechanisms governing polarization of cellular responses with ECM interactions.

Recent studies have indicated that PI3K and its downstream effector protein kinase B/Akt play important roles in regulating β-cell survival [13,48]. For example, once activated, Akt can inhibit several pro-apoptotic factors such as forkhead box protein O (FOXO), Bcl-2-associated death promoter (Bad), Bcl-2-associated X protein (Bax), and caspase-9 [13,49]. In several experimental models, active PKCδ has been shown to promote apoptosis by inhibiting Akt [50,51]. Our results further support a role for PKCδ in regulating Akt activity and in suppressing the PI3K/Akt pathway in response to pro-inflammatory cytokines, leading to β-cell apoptosis. Nevertheless, in the presence of cytokines pro-survival Akt signaling is only modestly increased compared to RTG alone or RTG+Lys, suggesting that laminin may also protect β-cells against cytokine-induced death through an Akt-independent pathway. In neutrophils for example, activation of β1 integrins prevents activation of mitogen activated protein kinase (MAPK) signaling to promote cell survival [52]. In islet β-cell, suppression of p38 MAPK signaling has been shown to promote survival in the INS1E rat β-cell line, suggesting that this alternative pathway may partially mediated laminin-mediated β-cell survival [53]. Additionally, as full activation of Akt requires phosphorylation at both Thr^308^ and Ser^473^ sites, future studies are warranted that examine the effect of laminin and PKCδ on phosphorylation of Akt at Ser^473^ in addition to Thr^308^ that was analyzed in this study to further confirm the role of Akt signaling [13]. Altogether, we show that laminin interactions with the β1 integrin on the surface of β-cells increases the activity of pro-survival Akt, likely by downregulating PKCδ and reducing levels of DAG in the β cell, conferring protection against cytokine induced death.

## Conclusion

In summary, we have identified a role for laminin in protecting β-cells against cytokine-induced death through activation of β1 integrin signaling leading to reduced DAG content and reduced activation of pro-apoptotic PKCδ signaling in MIN6 cells, mouse islets, and human islets. Our results also suggest that laminin induces downstream activation of pro-survival Akt that is partially mediated by PKCδ activity. Overall, our findings provide novel insight into the role of laminin in activating pro-survival signaling pathways through downregulation of the lipid second messenger DAG and reduced activation of PKCδ in pancreatic β-cells. Our results provide strong mechanistic evidence for the pancreatic islet microenvironment in regulating islet survival. Further, our results suggest that loss of laminin, as occurs during the progression of T1D, may render islet β-cells more susceptible to immune-mediated death, supporting laminin-β1 integrin signaling as a potential therapeutic target for preserving β-cell integrity during T1D.

## Materials & Methods

### 1. RTG synthesis

Poly(serinol hexamethylene urea)-co-poly(N-isopropylacylamide) (PSHU-PNIPAAm) was synthesized following a previously established protocol [29–31]. Briefly, 6 mmol N-BOC-serinol (1.147 g; Sigma-Aldrich, St. Louis, MO) and 6 mmol urea (0.36 g; Fisher Scientific, Hampton, NH) were dissolved in 6 mL N-N-Dimethylformamide (DMF; Sigma-Aldrich, St. Louis, MO) in a 90°C mineral oil bath with constant stirring. Under a nitrogen atmosphere, 12 mmol hexamethylene diisocyanate (HDI; 2.018 g; Sigma-Aldrich, St. Louis, MO) was added dropwise and allowed to react for 7 days protected from light. The reaction was precipitated in anhydrous diethyl ether (Fisher Scientific, Hampton, NH) and dried via rotary evaporation on a Rotavapor R-100 (BUCHI, New Castle, DE) to yield poly(serinol hexamethylene urea) (PSHU).

PSHU was deprotected with 30 mL of a 50:50 mixture containing dichloromethane (Sigma-Aldrich, St Louis, MO) and trifluoroacetic acid (TFA; Sigma-Aldrich, St. Louis, MO) for 45 min at room temperature. The acid was removed via rotary evaporation and the product was re-dissolved in DMF, then precipitated in anhydrous diethyl ether. The resulting polymer was redissolved in 2,2,2-trifluoroethylene (TFE; Sigma-Aldrich, St. Louis, MO) to yield de-protected PSHU (dPSHU).

Carboxylic acid-terminated PNIPAAm (PNIPAAm-COOH) was synthesized by reacting 800 mmol poly(N-isopropylacrylamide) (NIPAAm; 5 g; Sigma-Aldrich, St. Louis, MO) and 4 mmol 4,4-azobis (ACVA; 0.06 g; 4-cyanovaleric acid) (Sigma-Aldrich, St. Louis, MO) in 10 mL DMF under nitrogen atmosphere for 3 h at 68°C with constant stirring. The mixture was precipitated in 60°C water, then dissolved in room temperature water. The product was dialyzed for 48 h in 3500 Da molecular weight cut-off (MWCO) dialysis tubing (Fisher Scientific, Hampton, NH) and lyophilized to yield a white product.

Carbodiimide linking chemistry was used to conjugate PNIPAAm-COOH onto PSHU-NH_2_. PNIPAAm-COOH was dissolved in 5 mL anhydrous DMF and slowly added to a flask containing five molar excess of N-hydroxysuccinamide (NHS; Alfa Aesar, Ward Hill, MA) and N-ethyl-N’-(3-dimethylaminopropyl)carbodiimide hydrochloride (EDC-HCl; Sigma-Aldrich, St. Louis, MO). The reaction was performed under a nitrogen atmosphere and was protected from light for 24 h at room temperature. dPSHU was dissolved in 1 mL DMF and added dropwise to the conjugation reaction, and the reaction proceeded for 24 h under a nitrogen atmosphere. The mixture was precipitated in room temperature dH_2_O and dialyzed for 48 h in 12000 Da MWCO dialysis tubing (cat. no. 08–670-3A; Fisher Scientific, Hampton, NH) to remove unreacted PNIPAAm and lyophilized to yield the reverse thermal gel (RTG) PSHU-PNIPAAm.

### 2. Laminin-511 purification

Human embryonic kidney (HEK-293) cell lines stably expressing recombinant laminin-511 (obtained from Dr. Yurchenco Lab, Rutgers University) were cultured in 1X DMEM medium containing 4.5 g/L D-glucose and L-glutamine (Fisher Scientific, Hampton, NH) and supplemented with 10% fetal bovine serum (FBS; Fisher Scientific, Hampton, NH) and 5% penicillin/streptomycin (Sigma-Aldrich, St. Louis, MO) at 37°C and 5% CO_2_. After 5 days, culture medium was changed to 1X DMEM medium containing 1 μg/mL puromycin (Fisher Scientific, Hampton, NH), 100 μg/mL zeocin (Fisher Scientific, Hampton, NH), and 500 μg/mL geneticin (Fisher Scientific, Hampton, NH). Cells were cultured for 5 more days, and the media was collected for affinity purification. The collected media was centrifuged at 4000 rpm for 5 min to pellet the cells and obtain the supernatant. The collected supernatant was then filtered with a 0.2 μm filter and stored at −80°C.

Recombinant laminin was purified from the media using column chromatography on a heparin-agarose column (Sigma-Aldrich, St. Louis, MO) and a FLAG M2-agarose column (Sigma-Aldrich, St. Louis, MO) per the manufacturer’s instructions and as described [54]. Purified laminin was aliquoted and stored at −80°C, and western blot analysis was used to confirm the expression of laminin.

### 3. Synthesis of RTG+Lys

RTG+Lys was synthesized based on a previously described protocol [30]. Briefly, L-lysine monohydrochloride (Sigma-Aldrich, St. Louis, MO) was dissolved in 5 mL 1X PBS with five molar excess of EDC-HCl and NHS in a 25 mL round-bottom flask and the mixture was stirred for 15 min at 4°C. A 10 mL solution of 1 g PSHU-PNIPAAm dissolved in 10 mL 1X PBS was added dropwise, and the reaction was performed for 48 h at room temperature. The polymer was dialyzed (MWCO: 12000–14000 Da) for 5 days at room temperature, filtered through a 2 μm filter and lyophilized to yield RTG+Lys.

### 4. Synthesis of RTG+Lam

RTG+Lam was synthesized based on a previously described protocol [30]. Briefly, 2.5 × 10^−3^ mmol purified laminin-511 (1 mg) was dissolved in 5 mL 1X PBS with five molar excess of EDC-HCl and NHS in a 25 mL round-bottom flask and the mixture was stirred for 15 min at 4°C. A 10 mL solution of 1 g PSHU-PNIPAAm dissolved in 10 mL 1X PBS was added dropwise, and the reaction was performed for 48 h at 4°C. The polymer was dialyzed (MWCO: 12000–14000 Da) for 5 days at 4°C, filtered through a 2 μm filter, and lyophilized to yield RTG+Lam.

### 5. Nuclear magnetic resonance (NMR) spectroscopy

^1^H NMR spectra were collected on a 500 MHz liquid state NMR (JEOL USA, Inc., Peabody, MA). Spectra were collected in deuterated chloroform (DLM-7–100; Cambridge Isotope Laboratories, Inc., Andover, MA) and post-processed by means of Fourier Transform in iNMR (Nucleomatica).

### 6. Fourier-transform infrared spectroscopy (FTIR)

FTIR spectra were collected using NicoletiN 10MX infrared imaging microscope with the micro attenuated total reflectance (ATR) attachment.

### 7. Scanning electron microscopy (SEM)

5 wt% RTG, RTG+Lam, and RTG+Lys were prepared in RPMI medium, snap frozen in liquid nitrogen, and lyophilized for 24 h. Morphological characterization was carried out by scanning electron microscopy (SEM) using a Phenom tabletop SEM (ThermoFisher Scientific, Waltham, MA). The distance between RTG sheets was calculated by drawing straight lines between adjacent sheets and measuring the distance between them in ImageJ (NIH). Pore size was calculated as the average of two perpendicular diameters measured within the same pore.

### 8. Rheology

200 μL of 5 wt% RTG, RTG+Lam, RTG+Lys prepared in RPMI medium were loaded on a magnetic bearing rheometer (AR-G2, TA Instruments, New Castle, DE) between a 40 mm 2.0° cone and Peltier plate. Temperature sweep tests from 25°C to 40°C were conducted using a step size of 1°C at a constant frequency (1 Hz) and stress (0.05 Pa).

### 9. Animal care

All experiments using mice were performed at the University of Colorado Anschutz Medical Campus and in compliance with the guidelines and relevant laws set by the University of Colorado and the National Institutes of Health guide for the care and use of Laboratory animals. All experiments with mice were approved by the University of Colorado Institutional Animal Care and Use Committee (Protocol 00929). Mice were housed in a temperature- and light-controlled environment with 12-hour light-dark cycles, and they were provided access to food and drink ad libitum. Female C57Bl/6 mice were purchased from the Jackson Laboratories (strain #000664) at 8 weeks of age.

### 10. Human islets

Human islets were obtained from the Integrated Islet Distribution Program (IIDP) from the following donors as outlined in Table 1:

**Table 1:** Human cadaveric donor demographics and isolated islet viability and purity for islets obtained through the Integrated Islet Distribution Program (IIDP).

| RRID# | Viability | Purity | Age | Sex | BMI | Ethnicity/Race |
| --- | --- | --- | --- | --- | --- | --- |
| SAMN60580732 | 98% | 85% | 61 | Male | 30.16 | White |
| SAMN57330353 | 95% | 85% | 26 | Male | 25.10 | White |
| SAMN56769186 | 95% | 95% | 55 | Female | 26.40 | American Indian/Alaska Native |
| SAMN56446053 | 95% | 95% | 63 | Male | 33.50 | Black/African American |
| SAMN55907150 | 95% | 90% | 57 | Male | 27.90 | White |
| SAMN55343530 | 90% | 88% | 53 | Female | 34.30 | White |
| SAMN54553956 | 94% | 95% | 43 | Female | 26.50 | White |
| SAMN53125864 | 98% | 85% | 46 | Female | 40.70 | White |
| SAMN52936198 | 92% | 80% | 45 | Female | 26.60 | Hispanic/Latino |
| SAMN52617548 | 94% | 90% | 35 | Male | 26.50 | White |
| SAMN51198036 | 97% | 90% | 44 | Female | 31.00 | White |

Human donor tissue was de-identified through the IIDP and is therefore exempt from requiring human subject research protocols. Following delivery, human islets were cultured overnight in 1X 1640 RPMI medium with L-glutamine and 25 mM HEPES (Fisher Scientific, Hampton, NH) with 10% fetal bovine serum (FBS; Fisher Scientific, Hampton, NH), 10,000 U/mL penicillin and 10,000μg/mL streptomycin (Sigma-Aldrich, St. Louis, MO) before commencing experiments. Islets that were clear of any excess pancreatic tissue and were consistent in color and shape based on inspection through a microscope were selected for use in experiments.

### 11. Mouse islet isolation and culture

MIP CreER expressing PKCδ WT (control) and PKCδ-βKO mice were injected with 50mg/kg tamoxifen in corn oil once daily for five consecutive days to induce recombination. Islets were isolated from 8 to 12-week-old C57Bl/6 or 8 to 12-week-old age and sex matched Cre expressing WT control and PKCδ-βKO mice two weeks after the first tamoxifen injection. For islet isolation, animals were injected with 100 mg/kg ketamine and 8 mg/kg xylazine and euthanized via exsanguination. Islets were isolated by injecting the pancreas with 12.5mg/mL collagenase, removing the pancreas, and performing enzymatic digestion at 37°C. Mouse islets were handpicked into 1X 1640 RPMI medium with L-glutamine and 25 mM HEPES with 10% FBS, 10,000 U/mL penicillin and 10,000μg/mL and incubated at 37°C and 5% CO_2_ overnight.

### 12. Islet encapsulation and treatment

Mouse and human islets were encapsulated in RTG, RTG+Lam, or RTG+Lys or freely cultured (unencapsulated) and either untreated or treated with a pro-inflammatory cytokine cocktail containing mouse or human recombinant cytokines at 1X relative cytokine concentration (1X RCC; 10 ng/ml TNF-α (410-MT, 210-TA), 5 ng/ml IL-1β (401-ML, 201-LB), and 100 ng/ml IFN-γ (485-MI, 285-IF), R&D Systems, Minneapolis, MN) respectively for 24 h. As a positive control for β-cell integrin interactions, islets were treated with 1 μM of the β1 integrin agonist pyrintegrin (cat. no. HY-13306; MedChemExpress, Monmouth Junction, NJ) for 24 h. As a negative control for β-cell integrin interactions, islets were pre-treated with 100 ng/mL of the β1 integrin antagonist AIIB2 (RRID: AB_528306; Developmental Studies Hybridoma Bank, Iowa City, IA) for 30 min prior to encapsulation and during culture in RTG.

### 13. MIN6 cell culture

MIN6 cells were cultured in 1X DMEM medium containing 4.5 g/L D-glucose and L-glutamine (Fisher Scientific, Hampton, NH) with 10% FBS (Fisher Scientific, Hampton, NH) and 5% penicillin/streptomycin at 37°C and 5% CO_2_. For experimental use, cells were passaged at 90% confluency and trypsinized with 0.05% Trypsin-EDTA (Fisher Scientific, Hampton, NH). Cell count and viability were noted prior to experimental use. For 2D culture, MIN6 cells ranged from passage 23-30 and were seeded at a density of ∼ 30,000 cells/cm^2^ in a 35mm dish.

### 14. Immunohistochemistry

MIN6 cells, C57Bl/6 mouse islets and human islets were fixed with 4% paraformaldehyde (Fisher Scientific, Hampton, NH) in PBS for 10 min at 37°C. The islets were then permeabilized in PBS with 5% normal donkey serum (Sigma-Aldrich, St. Louis, MO) for 5 min at 37°C. After permeabilization, the islets were incubated in primary antibody solution containing mouse anti-insulin monoclonal antibody, Alexa Fluor 488 (RRID: AB_2574469, cat. no. 53-9769-82, Fisher Scientific, Hampton, NH) diluted 1:100, rat anti-β1 integrin antibody (AIIB2; RRID: AB_528306, Developmental Studies Hybridoma Bank, Iowa City, IA) diluted 1:4, and rabbit anti-glucagon antibody (cat. no. 2760, Cell Signaling Technology, Danvers, MA) diluted 1:200 in PBS with 5% normal donkey serum for 1 h at 37°C. After incubation in the primary antibody solution, the islets were washed three times with PBS and incubated in a secondary antibody solution containing goat anti-rat secondary antibody, Alexa Fluor 647 (RRID: AB_141778, cat. no. A-21247, Fisher Scientific, Hampton, NH) diluted 1:200 in PBS with 5% normal donkey serum for 1 h at 37°C. Islets were washed three times before mounting in DAPI-Fluoromount-G^TM^ (Electron Microscopy Sciences, Morgantown, PA) for at least 30 min at 37°C, then imaged.

C57Bl/6 mouse islets encapsulated in RTG+Lam were fixed and permeabilized as above and antigen retrieval was performed in 1X citrate buffer, pH 6 (Sigma-Aldrich, St. Louis, MO) at 60°C for 2 h. After antigen retrieval, the encapsulated islets were incubated in primary antibody solution containing mouse anti-insulin monoclonal antibody, Alexa Fluor 488 (RRID: AB_2574469, cat. no. 53-9769-82, Fisher Scientific, Hampton, NH) diluted 1:100 and rabbit anti-laminin antibody (cat. no. ab11575, Abcam, Waltham, MA) diluted 1:250. After incubation in the primary antibody solution, the islets were washed three times with PBS and incubated in a secondary antibody solution containing goat anti-rabbit secondary antibody, Alexa Fluor 568 (RRID: AB_143157, cat. no. A-11011, Fisher Scientific, Hampton, NH).

### 15. Islet viability

Islets were incubated with 15 μL/mL propidium iodide (Sigma-Aldrich, St. Louis, MO) and 15 μL/mL fluorescein diacetate (Sigma-Aldrich, St. Louis, MO) in BMHH imaging buffer (125 mM NaCl, 5.7 mM KCl, 2.5 mM CaCl_2_, 1.2 mM MgCl_2_, and 10 mM HEPES in dH2O, 0.1% bovine serum albumin, pH 7.4) for 10 min. Imaging was performed on a Leica Stellaris 5 confocal microscope with a 40X water immersion objective. 488 nm and 514 nm solid state lasers were used for excitation, and emission was collected with HyD spectral detectors. All islets were imaged as a z-stack consisting of three images 8-10 μm apart and live/dead cells were counted manually in ImageJ (NIH).

### 16. Förster resonance energy transfer (FRET)

PKCδ activity at the cell membrane was determined using a FRET-based activity sensor (δCKAR) containing a membrane localization sequence, which was generously given by Dr. Alexandra Newton and previously described [33]. The δCKAR construct was transfected into MIN6 cells with Lipofectamine 3000 (Thermo Fisher Scientific) per the manufacturer’s instructions. Transfected cells were encapsulated in RTG, RTG+Lam, or RTG+Lys for 24 h and imaged on a Leica Stellaris confocal microscope with a 40x water immersion objective. 448 nm and 514 nm solid state lasers were used for excitation, and emission was collected with HyD spectral detectors. Cells were imaged for 5 min under basal conditions, then for 25 min after treatment with the cytokine cocktail (10 ng/mL TNF-α, 5 ng/mL IL-1β, and 100 ng/mL IFN-γ). As a positive control, cells were subsequently treated with 300 nM of the PKCδ-specific activator phorbol 12-myristate 13-acetate (PMA, Sigma-Aldrich, St. Louis, MO) and imaged for an additional 5 minutes. FRET images were obtained every 15 s. FRET efficiency was calculated in ImageJ using the equation

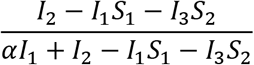

where I_1_, I_2_, and I_3_ correspond to donor, FRET, and acceptor channel intensities, respectively, and S_1_, S_2_ and α are correction factors for donor bleed-through, acceptor direct excitation, and donor crosstalk into the acceptor channel.

### 17. Western blotting

MIN6 cells were encapsulated in RTG, RTG+Lam, or RTG+Lys or freely cultured (unencapsulated) and either untreated or treated with a 1X pro-inflammatory cytokine cocktail (10 ng/ml TNF-α, 5 ng/ml IL-1β, and 100 ng/ml IFN-γ) for 1 h and 24 h. Each encapsulated sample contained ∼50,000 MIN6 cells in 200 μL RTG, RTG+Lam, or RTG+Lys and each unencapsulated sample contained ∼50,000 MIN6 cells in DMEM medium. Unencapsulated and encapsulated mouse and human islets were cultured for 24 h with treatments as previously described, using mouse or human recombinant cytokines, respectively. Each encapsulated sample contained 100 islets in 200 μL RTG, RTG+Lam, or RTG+Lys and each unencapsulated sample contained 100 islets in RPMI media. Islets and MIN6 cells were washed twice in 1X PBS and lysed by sonication for 30 s in lysis buffer containing 100 mM NaCl, 50 mM Tris-HCl, 10 mM MgCl_2_ with a mixture of protease inhibitors (ThermoFisher Scientific, Waltham, MA). Protein content was measured using the Pierce BCA Protein Assay Kit (cat. no. 23225; Fisher Scientific, Hampton, NH) per the manufacturer’s instructions. Samples were run on 4–15% mini-PROTEAN® TGX protein gels (Bio-Rad, Hercules, CA) and transferred to PVDF membranes (Azure Biosystems, Dublin, CA). The PVDF membranes were blocked in chemi-blot blocking buffer (Azure Biosystems, Dublin, CA) for 2 h at room temperature and probed with the following antibodies for >16 h at 4°C: rabbit anti-phospho-FAK (Tyr^397^; cat. no. 3283, Cell Signaling Technology, Danvers, MA) diluted 1:1000, rabbit anti-FAK (cat. no. 3285, Cell Signaling Technology, Danvers, MA) diluted 1:1000, rabbit anti-phospho-Akt (Thr^308^; cat. no. 13038, Cell Signaling Technology, Danvers, MA) diluted 1:1000, rabbit anti-Akt (pan; cat. no. 4691, Cell Signaling Technology, Danvers, MA) diluted 1:1000, rabbit anti-PKCδ (cat. no. ab182126, Abcam, Waltham, MA) diluted 1:5000 and mouse anti-β-actin (cat. no. sc-47778, Santa Cruz Biotechnology, Santa Cruz, CA) diluted 1:100. All membranes were stained with either goat anti-rabbit horseradish peroxidase (HRP)-conjugated secondary antibody (cat. no. 102649-670; VWR) diluted 1:10,000 or goat anti-mouse HRP-conjugated secondary antibody (RRID: AB_2533947, cat. no. 62-6520, Fisher Scientific, Hampton, NH) diluted 1:1000 for 2 h at room temperature. The membranes were incubated with Radiance Plus (Azure Biosystems, Dublin, CA) for 2 min in the dark and then visualized using the Azure c600 imaging system (Azure Biosystems, Dublin, CA). Protein quantification was performed with ImageJ (NIH) using densitometric analysis.

Purified laminin fractions and TBS buffer (50 mM Tris pH 7.4, 150 mM NaCl, 1 mM EDTA, 0.05% NaN_3_) controls were blocked and probed with mouse anti-laminin-5 antibody (cat. no. sc-13586, Santa Cruz Biotechnology, Santa Cruz, CA) diluted 1:200, followed by goat anti-mouse HRP-conjugated secondary antibody (RRID: AB_2533947, cat. no. 62-6520, Fisher Scientific, Hampton, NH) diluted 1:1000. Protein concentration of purified laminin fractions was determined by BCA assay prior to loading, and buffer controls were loaded at an equivalent volume.

### 18. Diacylglycerol (DAG) assay

MIN6 cells were grown on 75 cm^2^ tissue culture-treated flasks that were either uncoated or coated with full-length human recombinant laminin-511 (Biolamina, Sundbyberg, Sweden) until 90% confluency. After confluency, the cells were either untreated or treated with 1X cytokines in DMEM medium for 24 h. Then, the media was removed, and the cells were washed twice with cold PBS. The cells were harvested using a rubber policeman and each sample contained roughly 1×10^7^ cells that were used for the assay. DAG content was measured using the DAG Assay Kit (cat. no. MET-5028, Cell Biolabs, San Diego, CA) according to the manufacturer’s instructions.

### 19. Statistical analysis

Statistical analysis was performed using Origin software (OriginLabs, Northampton, MA). One sample t-test and one-way ANOVA with Tukey’s post hoc analysis were performed as indicated. A p-value of <0.05 was considered statistically significant.

## Author contribution statement

All authors made substantial contributions to study design, data acquisition and analysis, and manuscript revision. C.E. designed experiments, led data collection and wrote manuscript. M.H.S.M. and K.H. designed experiments and performed data collection and analysis. G.G. and R.S. performed data collection and analysis.

N.L.F. designed experiments, acquired funding, supervised, and edited the manuscript. D.P. and B.P provided resources and experimental guidance and edited the manuscript. R.K.P.B. acquired funding, provided experimental guidance, and edited the manuscript. All authors have read and approved the final manuscript for submission.

