## Supplemental Figures for "Laminin-511 protects pancreatic β-cells from cytokine-induced death through integrin-mediated pro-survival signaling and modulation of protein kinase C δ"

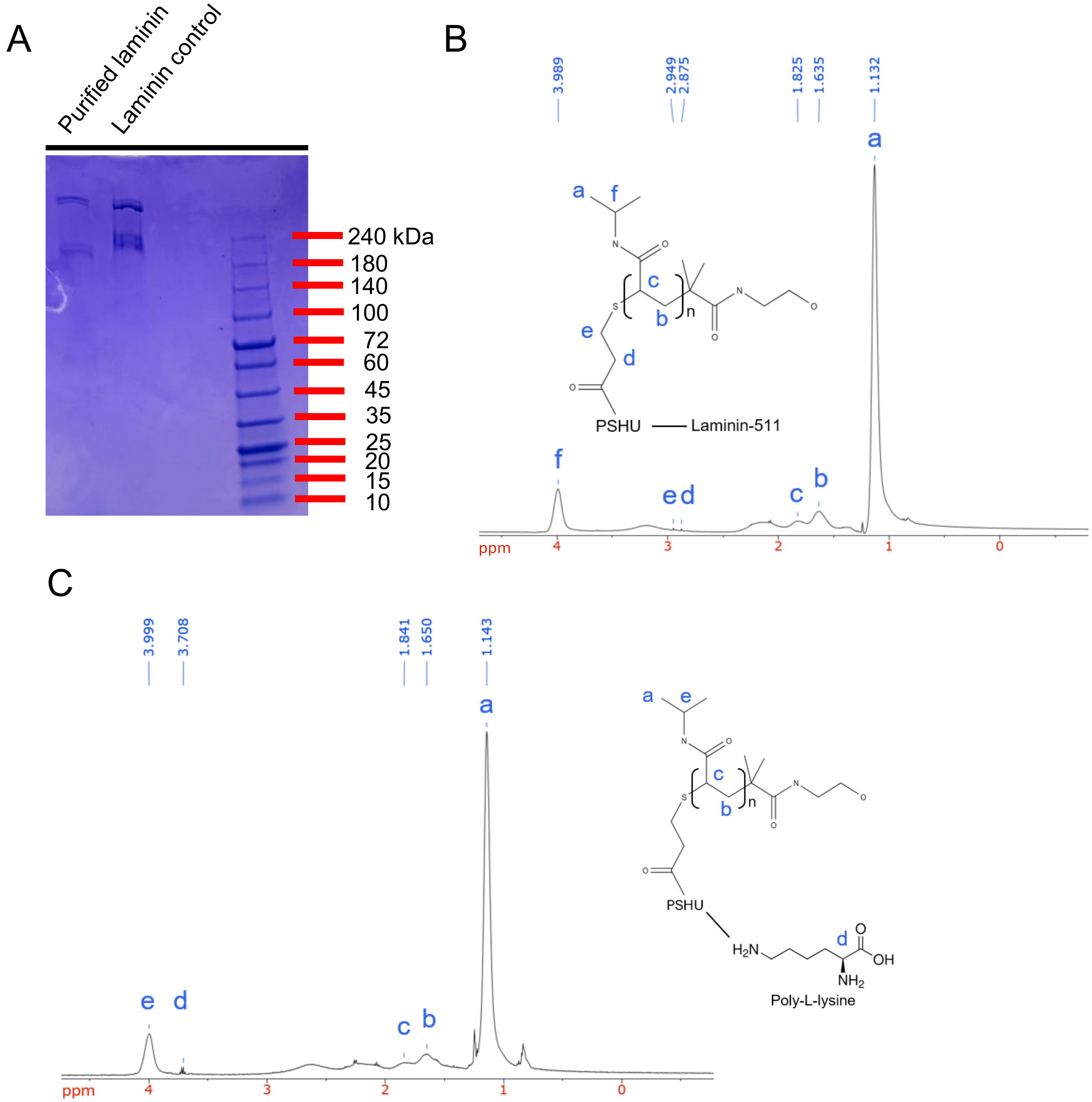


**Supplemental Figure 1: Chemical characterization of RTG+Lam and RTG+Lys.** (A) Representative polyacrylamide gel stained with Coomassie Brilliant Blue showing purified recombinant laminin-511 isolated from HEK293 cell conditioned medium. Commercial recombinant laminin-511 (BioLamina) was included as a positive control. (B) $^{1}H$ NMR spectrum of PSHU-PNIPAAm-laminin. Characteristic PNIPAAm peaks are observed at 1.13 and 1.64 ppm, corresponding to (a) methyl and (b) methylene protons, respectively. Additional peaks (c) and (f) at 1.74 and 3.97 ppm are consistent with the PNIPAAm backbone methine proton and the isopropyl methine proton, respectively. Peaks (d) and (e) at 1.83 and 3.99 ppm are consistent with methylene protons adjacent to sulfur- and carbonyl-containing groups. (C) $^{1}H$ NMR spectrum of PSHU-PNIPAAm+poly-L-lysine. Characteristic PNIPAAm peaks are observed at 1.14 and 1.65 ppm, corresponding to (a) methyl and (b) methylene protons, respectively. Additional peaks (c) and (e) at 1.84 and 3.99 ppm are consistent with the PNIPAAm backbone methine proton and the isopropyl methine proton, respectively. Peak (d) is consistent with the α-proton of poly-L-lysine.

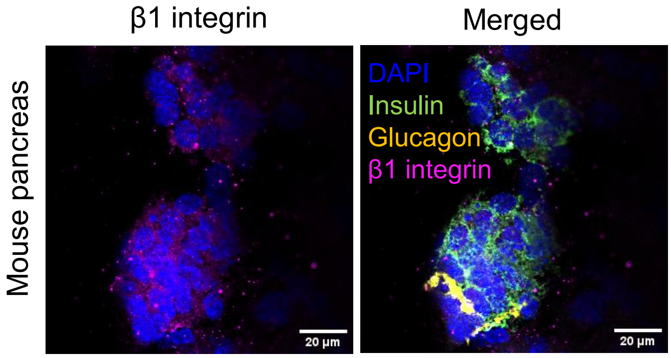


**Supplemental Figure 2: Distribution of laminin and β1 integrins around islet cells.** IHC images of a B6 mouse pancreas slice stained with DAPI (blue), insulin (green), glucagon (yellow) and β1 integrin (magenta).

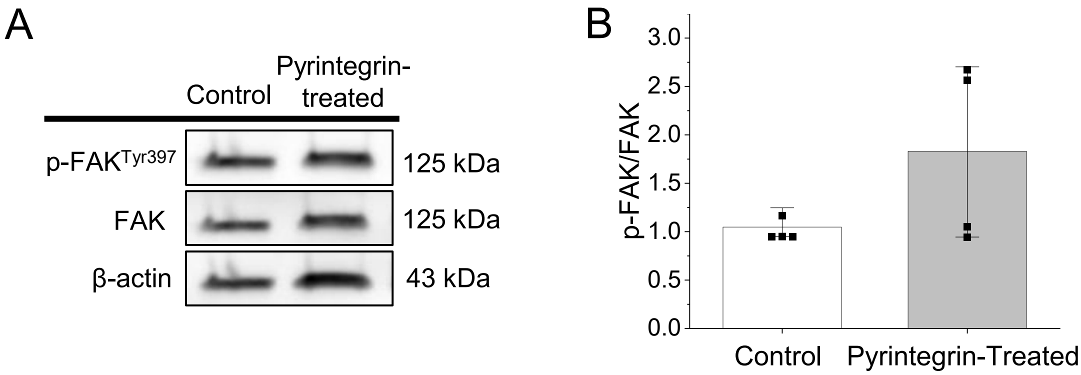


**Supplemental Figure 3: Pyrintegrin promotes FAK activation in mouse islets.** (A) Representative western blot image of p-FAK, FAK and β actin present in mouse islets untreated or treated with the β1 integrin agonist pyrintegrin for 24 h. (B) Western blot quantification for p-FAK/FAK in mouse islets untreated or treated with pyrintegrin (n=4). Error bars represent the mean +/- SEM. p-value <0.05 is significant as determined by ANOVA.

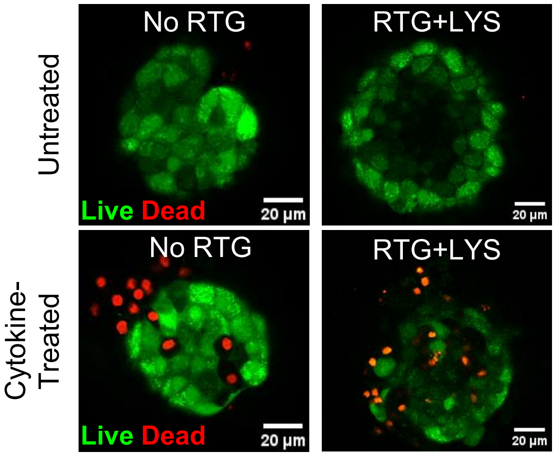


**Supplemental Figure 4: Laminin improves mouse islet viability under cytokine stress.** (A) Representative images of mouse islet viability in no RTG and 5 wt% RTG+Lys either untreated or treated with cytokines for 24 h.

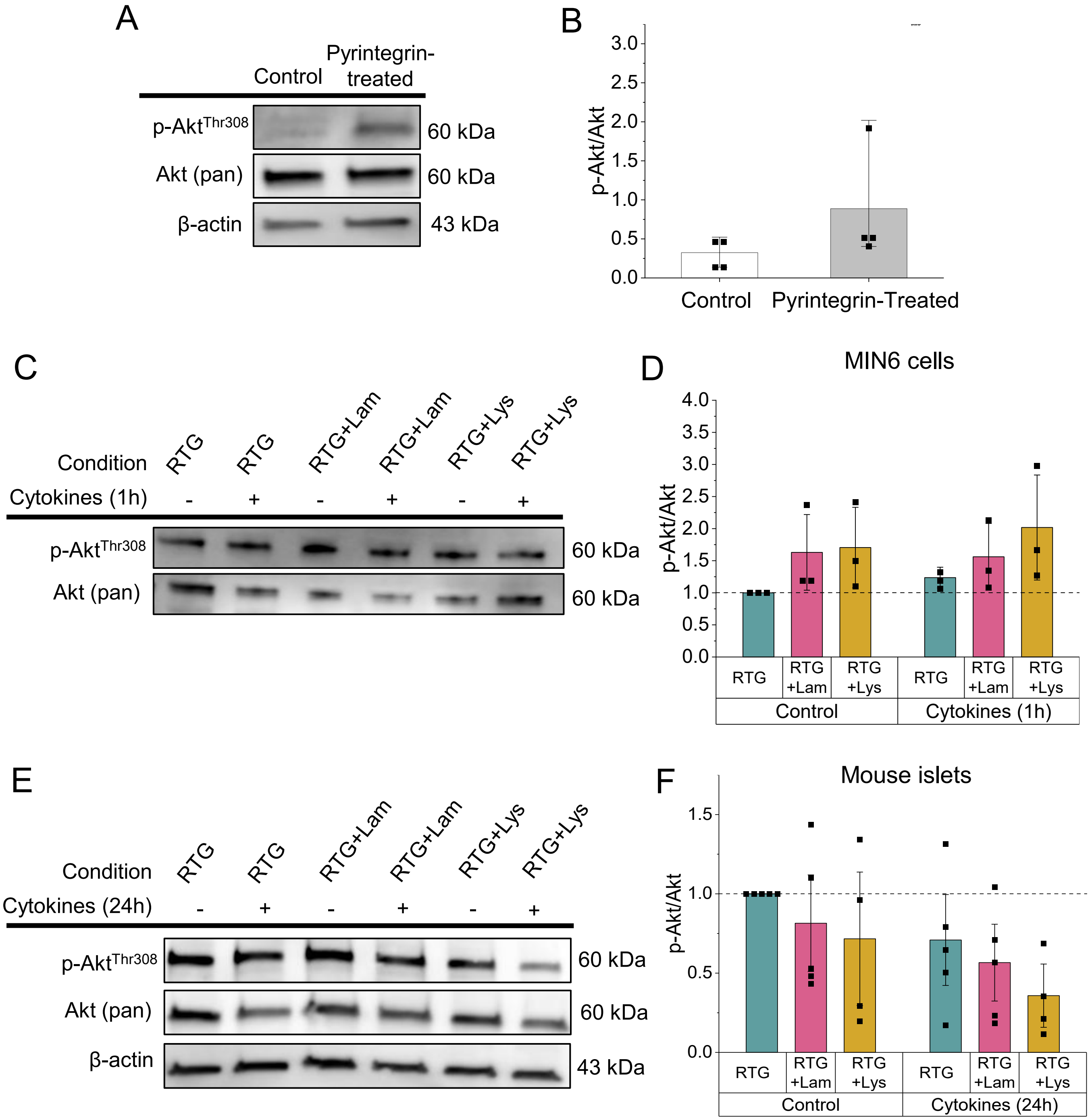


**Supplemental Figure 5: Akt signaling in encapsulated MIN6 cells and mouse islets.** (A) Representative western blot image of p-Akt, Akt and β actin present in mouse islets untreated or treated with the β1 integrin agonist pyrintegrin for 24 h. (B) Western blot quantification for p-Akt/Akt in mouse islets untreated or treated with pyrintegrin (n=4). (C) Representative western blot image of p-Akt and Akt present in MIN6 cells encapsulated in RTG, RTG+Lam, or RTG+Lys for 1 h and treated with cytokines for 1 h. (D) Western blot quantification for p-Akt/Akt in MIN6 cells (n=3). Data is normalized to RTG (control). (E) Representative western blot image of p-Akt, Akt and β actin present in mouse islets encapsulated in RTG, RTG+Lam, or RTG+Lys and treated with cytokines for 24 h. (F) Western blot quantification for p-Akt/Akt in mouse islets encapsulated in RTG untreated (n=5), RTG-Lam untreated (n=5), RTG-Lys untreated (n=4), RTG + cytokines (n=5), RTG+Lam +cytokines (n=5), or RTG+Lys + cytokines (n=4) for 24 h. Data is normalized to RTG (control). Error bars represent the mean +/- SEM. Differences among treatment groups were assessed separately within the control and cytokine-treated conditions using one-way ANOVA. Statistical significance was defined as $p < 0.05$.
